# Unbiased metagenomic sequencing for pediatric meningitis in Bangladesh reveals neuroinvasive Chikungunya virus outbreak and other unrealized pathogens

**DOI:** 10.1101/579532

**Authors:** Senjuti Saha, Akshaya Ramesh, Katrina Kalantar, Roly Malaker, Md Hasanuzzaman, Lillian M. Khan, Madeline Y Mayday, M S I Sajib, Lucy M. Li, Charles Langelier, Hafizur Rahman, Emily D. Crawford, Cristina M. Tato, Maksuda Islam, Yun-Fang Juan, Charles de Bourcy, Boris Dimitrov, James Wang, Jennifer Tang, Jonathan Sheu, Rebecca Egger, Tiago Rodrigues De Carvalho, Michael R. Wilson, Samir K Saha, Joseph L DeRisi

**Author notes:** Corresponding author Senjuti Saha Child Health Research Foundation, Department of Microbiology, Dhaka Shishu Hospital, Sher-e-Bangla Nagar, Dhaka, Bangladesh. Co-corresponding author Joseph L DeRisi Department of Biochemistry and Biophysics, 700 4th St, Byers Hall s403c, University of California, San Francisco, California, United States of America.

## Abstract

The disease burden due to meningitis in low and middle-income countries remains significant and failure to determine an etiology impedes appropriate treatment for patients and evidence-based policy decisions for populations. Broad-range pathogen surveillance using metagenomic next-generation sequencing (mNGS) of RNA isolated from cerebral spinal fluid (CSF) provides an unbiased assessment for possible infectious etiologies. In this study, our objective was to use mNGS to identify etiologies of pediatric meningitis in Bangladesh.

We conducted a retrospective case-control mNGS study on CSF from patients with known neurologic infections (n=36), idiopathic meningitis (n=25), without infection (n=30) and six environmental samples collected between 2012-2018. Using an open-access, cloud-based bioinformatics pipeline (IDseq) and machine learning, we identified potential pathogens which were confirmed through qPCR and Sanger sequencing. These cases were followed-up through phone/home-visits. The CSF samples were collected from children with WHO-defined meningeal signs during prospective meningitis surveillance at the largest pediatric referral hospital in Bangladesh.

The 91 participants (42% female) ranged in age from 0-160 months (median: 9 months). In samples with known infectious causes of meningitis and without infections (n=66), there was 83% concordance between mNGS and conventional testing. In idiopathic cases (n=25), mNGS identified a potential etiology in 40% (n=10), including bacterial and viral pathogens. There were three instances of neuroinvasive Chikungunya virus (CHIKV). The CHIKV genomes were >99% identical to each other and to a Bangladeshi strain only previously recognized to cause systemic illness in 2017. CHIKV qPCR of all remaining stored CSF samples from children who presented with idiopathic meningitis in 2017 at the same hospital (n=472) revealed 17 additional CHIKV meningitis cases. Orthogonal molecular confirmation of each mNGS-identified infection, case-based clinical data, and follow-up of patients substantiated the key findings.

Using mNGS, we obtained a microbiological diagnosis for 40% of idiopathic meningitis cases and identified a previous unappreciated pediatric CHIKV meningitis outbreak. Case-control CSF mNGS surveys can complement conventional diagnostic methods to identify etiologies of meningitis and facilitate informed policy decisions.

## Introduction

Globally there are 10.6 million cases of meningitis and 288,000 deaths every year.^1,2^ In addition, at least a quarter of survivors suffer from long-term neurological sequelae.^3^ The vast majority of meningitis cases occur in low- and middle-income countries (LMICs).^4^ In a World Health Organization (WHO)-supported meningitis surveillance study in Dhaka, Bangladesh,^5^ we collected 23,140 cerebrospinal fluid (CSF) samples from patients with suspected meningitis between 2004 and 2016, 8,125 of which contained ≥10 WBC/µl. We were able to detect a bacterial etiology in only 1,585 (20%) of these cases despite the use of multiple diagnostic tools including culture, serologic and antigen assays and pathogen-specific qPCR. Such low rates of microbiological diagnosis are common in many settings globally, hampering implementation of evidence-based policy decisions for optimizing local empiric treatment protocols and disease prevention strategies.^6,7^

The challenges of obtaining a microbiological diagnosis may be due to a combination of multiple factors including (i) meningitis is caused by a wide variety of microbes, some of which are uncommon and lack diagnostic assays, (ii) prior antibiotic exposure and delay in care-seeking can lower the yield of culture and PCR-based methods, and/or (iii) non-infectious causes of inflammation can mimic infectious meningitis. Drawing on recent studies demonstrating the promise of unbiased metagenomic next-generation sequencing (mNGS) approaches to identify pathogens in diverse biological specimens,^8–11^ we sought to conduct a retrospective case-control study to investigate CSF of children with idiopathic meningitis in Bangladesh.

## Methods

### Study site and population

All CSF samples used in this study were collected as part of the meningitis surveillance study supported by the WHO conducted in Dhaka Shishu Hospital (DSH). Children admitted at DSH were enrolled if they met WHO-defined inclusion criteria of meningitis and if a CSF specimen was collected (Table S1).^12^

Protocols were approved by the ethical review board of the Bangladesh Institute of Child Health. Samples were collected for routine clinical care, at the discretion of the attending physician. Informed consent was obtained from parents/caregivers.

### Laboratory methods and data collection

CSF specimens were cultured (Figure 1A) using the standard procedure and pneumococcal antigen was detected by immunochromatographic test (BinaxNow).^13–15^ White blood cells (WBC) in the specimens were counted and differentiated into lymphocytes and polymorphonuclear neutrophils (PMNs). Culture-negative and pneumococcus-antigen-negative CSF specimens underwent latex agglutination and PCR testing for *Haemophilus influenzae*, pneumococcus and meningococcus, the predominant bacterial etiologies of meningitis in the region. Surplus CSF was stored at −80°C. Detection of chikungunya virus (CHIKV) was conducted using qPCR with published primers on all CSF specimens stored and collected in 2017 (n=472) (Table S2).^16^

**Figure 1.**
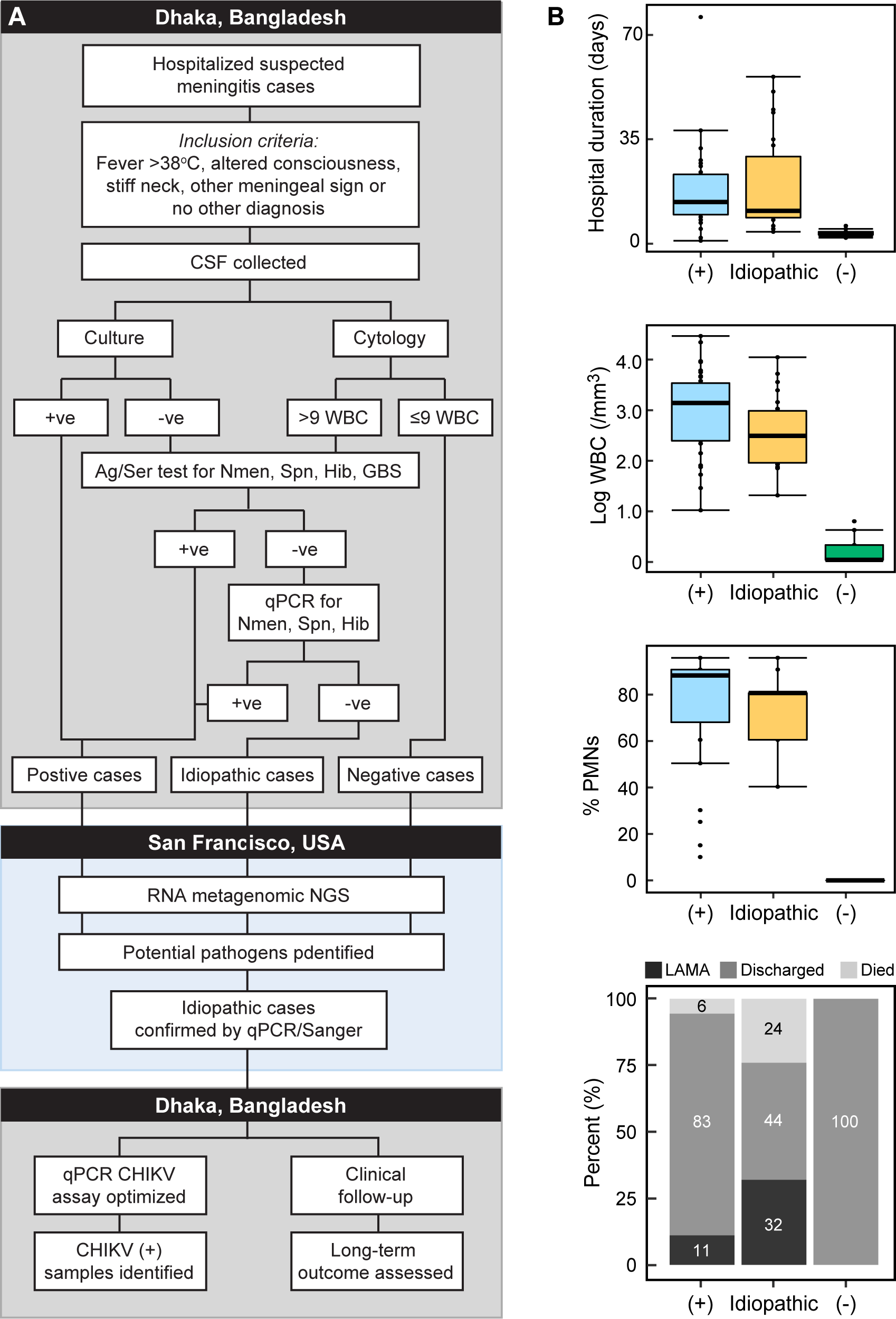
Selection and characteristics of samp les used in this study. A. Study flow diagram. B. Comparison of clinical characteristics between the three types of samples, positive (+), idiopathic, and negative (-) chosen for this study. Ag: antigen; Ser: serology; Spn: *Streptococcus pneumoniae*; Nmen: *Neisseria meningitidis*; Hib: *Haemophilus influenzae* tybe b; GBS: Group B Streptococcus; CHIKV: Chikungunya virus; PMN: Polymorphonuclear neutrophils

### Sample selection

Samples collected between 2012 and 2018 were selected and clinical details of all selected samples are provided in Table S3. For positive controls, CSF specimens where an etiology could be successfully established through culture, serology, antigen testing, and/or qPCR were chosen. For idiopathic samples, specimens were randomly chosen from a set that contained ≥20 WBC/µl (≥40% PMNs) (median: 314 WBC/µl), and ≥40 mg/dl protein (median: 220 mg/dL) (normal range is 15-45 mg/dL).

Negative controls consisted of randomly chosen CSF specimens from patients in whom an alternate diagnosis was ultimately made, the child was discharged within 6 days of hospitalization, and CSF samples contained ≤6 WBC/µl (median: 0 WBC/µl) and ≤30 mg/dl protein (median: 20 mg/dL). This set also included environmental samples, which were nuclease-free water (Invitrogen, 10977-015) samples (n=5) transferred into CSF collection tubes in the patient wards and treated and stored in the laboratory like CSF specimens in a blinded fashion. A “no template” water control sample was included during RNA extraction.

### mNGS and confirmatory testing

Total RNA was extracted from 100 µl of unspun CSF, and mNGS libraries were prepared following published methods.^17^ External RNA Controls 103 Consortium collection (ERCC) (ThermoFisher, 4456740) spike-in controls were used in every sample as markers of potential library preparation errors and to permit calculation of input RNA mass. Libraries were sequenced on a NovaSeq 6000 to generate 150 bp, paired-end sequences. All non-human sequence reads were deposited in Sequence Read Archive, National Center for Biotechnology Information (NCBI) BioProject PRJNA516582.

Pathogens were identified from the raw fastq files using the IDseq portal, a cloud-based, open-source bioinformatics platform designed for detection of microbes from metagenomic data (Figure S1, Methods S1). Similar to published methods, potentially pathogenic microbes were distinguished from both ubiquitous environmental contaminants and commensal flora using a Z-score metric for each genus relative to a background distribution derived from the set of CSF specimens from non-meningitis cases and water controls.^8^ Taxa with a Z-score less than one were removed from analysis. To further aid analysis, we employed a published logistic regression method to classify and assign potential etiological candidates in each sample.^18^ We retrained the model using the following features: RNA-seq reads per million (rpM), rank amongst all detected microbes within the sample, and a binary variable indicating whether the microbe has established pathogenicity.^18,19^ Microbes with probability scores >0.2 were reported as potential pathogens. In cases where more than one potential pathogen was identified, only the top scoring pathogen was considered. Based on the water controls, a minimum calculated RNA input threshold of 3.0 pg was required for pathogen prediction. The average RNA input of the set of non-infectious CSF samples was 1.6 pg (range: 0.9–2.8).

Potential pathogens identified by mNGS were confirmed through PCR and Sanger sequencing (Table S2).

### Genome assembly, microbial typing, phylogenetics

For *de novo* assembly and annotation of draft genomes, we used the St. Petersburg genome assembler (SPAdes, v3.11.1)^20^ and Geneious (v10.3.2).^21^ Genotype assignments for viruses were identified using BLASTn, with the assembled genome sequence of the virus as query. We specifically compared the assembled CHIKV genomes with selected CHIKV genomes available in NCBI data for time-resolved phylogenetics (Method S2).

### Clinical Data collection and patient follow-up

Clinical and demographic data were collected from electronically stored surveillance forms. The number and distribution of suspected and confirmed CHIKV febrile cases were collected from the microbiology laboratory records of Shishu Shasthya Foundation Hospital and DSH, the two largest pediatric hospitals of Bangladesh, serving the same catchment area. Clinical follow-up was conducted through telephone and/or home visits using structured questionnaires.

## Results

CSF specimens were tested from 91 patients (42% female), ranging in age from 0-160 months (median: 9 months). Comparative clinical characteristics of the three case types: infectious, idiopathic and non-infectious, are provided in Figure 1B and Table S3. The majority (71%, n=65) of patients originated in Dhaka division, while the remaining cases sought care from 11 outlying districts (Figure S2A, B).

mNGS libraries (n=97) were prepared and sequenced (Methods), resulting in an average depth of 72 million reads/sample (IQR: 53-86 million). The resulting fastq files were processed using the open-source IDseq (www.IDseq.net, v1.8) platform (Figure S1, Methods S1). Using ERCC spike-in control RNAs (Methods), the calculated RNA input masses were highly correlated with WBC count (log scaled Pearson r=0.69).^17^ RNA input masses for the non-infectious CSF samples and template-free controls were significantly less than both the samples of known and unknown etiology (2 pg vs 273 pg, p <0.0001 by Wilcoxon rank sums) (Fig 2, Table S4).

**Figure 2.**
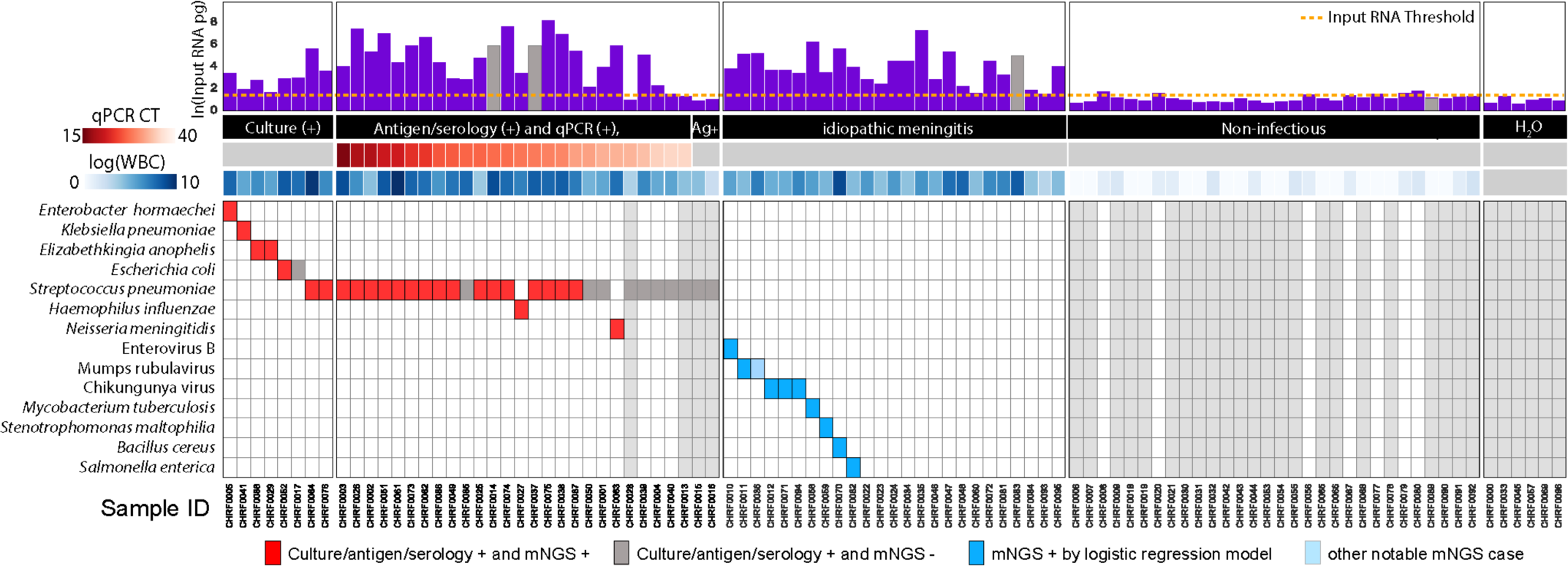
Pathogen Identification through mNGS and logistic regression in all sample types. Total input RNA (log pg) is shown for all samples. Samples for which the input RNA values could not be reliably calculated (outliers), are shown as grey bars with imputed input RNA values based on the mean value in their respective groups (known infection, no infection, idiopathic samples). Samples in the known infection group are ordered by increasing cycle threshold, depicted as a heat map below the x-axis. Next, the WBC counts obtained by the clinical lab are also plotted as a heatmap. The predicted pathogens for all samples are shown as filled-in squares. Grey squares indicate samples which were considered positive by clinical diagnostic, but for whom no pathogen was detected by the pathogen-calling algorithm using mNGS data. Red boxes indicate concordant findings and blue boxes indicate new putative pathogens identified by mNGS data that were not identified by standard clinical methods. The light blue squares indicate pathogens that were not picked up by the logistic regression method but were flagged as potentially interesting by manual review and followed-up as if detected. Ag+: Antigen positive

### Specimens of known etiology (n = 36)

Positive control samples were drawn from cases with previously identified pathogens, via a combination of standard lab diagnostics, including culture (n=8), qPCR and antigen/serology (n=26), and antigen-only (n=2). Using logistic regression (Methods), we correctly identified pathogens in 7 of 8 (88%) samples that were culture positive (Figure 2B). For specimens that were previously PCR and antigen or serology positive, mNGS identified 24 of 27 (89%) samples whose confirmatory qPCR had a cycle threshold (Ct) of <32. Taking into account all specimens that were culture, PCR and/or antigen or serology positive (n=36) regardless of Ct value, 25 (69%) specimens were classified as containing potential pathogens by mNGS (Figure 2B).

### Negative specimens (n = 36)

Among the non-infectious specimens, only 4 (11%) had an input RNA mass greater than 3.0 pg. No potential pathogens were identified in these samples.

### Idiopathic specimens (n = 25)

Potential pathogens were identified in 10 of 25 (40%) idiopathic cases: four cases with bacterial pathogens (*Salmonella enterica, Stenotrophomonas maltophilia, Bacillus cereus, Mycobacterium tuberculosis)* and six with viral pathogens (CHIKV (n=3), mumps virus (n=2), enterovirus B) (Figure 2). All these were further confirmed with orthogonal PCR testing and Sanger sequencing investigation of case histories and patient follow-up.

### Salmonella enterica

CHRF0082: A 4-month old boy was admitted with suspected meningitis, with fever and bulging fontanel. The CSF specimen tested in this study was collected 8 days after admission and contained 400 WBC/µl (80% PMNs) and 450 µg/dl protein. The child was treated with ceftriaxone, netilmicin and amoxicillin, and discharged after 23 days with residual Bell’s palsy. mNGS identified *S. enterica* (Table S4), and retrospective investigation revealed that a CSF specimen was also collected the day of admission and that first specimen was culture positive for *S. enterica.* The child was healthy upon follow-up at 15 months of age.

### Mycobacterium tuberculosis

CHRF0058: A 13-year old girl was admitted after 30 days of fever, vomiting, and headache with neuroimaging observations of intracranial space occupying lesions. Her CSF had 360 WBC/µl (80% PMNs) and 200 µg/dl protein. After 15 days of treatment with empiric ceftriaxone, meropenem, azithromycin, and acyclovir, the family left against medical advice. mNGS identified *Mycobacterium tuberculosis* (TB). Follow-up revealed that the child went to several health care facilities, where she was ultimately diagnosed with TB meningitis based on clinical suspicion and initiated on anti-tubercular chemotherapy. The grandfather of the child lived in the same household and died from pulmonary TB 2-3 months before the onset of her symptoms. The child, after almost a year, remains bedridden with persistent neurocognitive impairment.

### Stenotrophomonas maltophilia

CHRF0059: A 4-month old girl was admitted for 10-days of fever, convulsion and cough. She was treated at home with empiric cefixime and azithromycin before seeking care at DSH. Her CSF sample contained 120 WBC/µl (80% PMNs) and 300 µg/dl protein. She was treated with ceftriaxone, meropenem, vancomycin, and amikacin and discharged after 28 days. *S. maltophilia* was detected by mNGS (Table S4). This child was lost to follow-up.

### Bacillus cereus

CHRF0070: A 6-day old boy was admitted with fever, convulsion, lethargy and yellow coloration of skin; the treating physicians provisionally diagnosed him with sepsis and neonatal jaundice. His CSF contained 12,000 WBC/µl (95% PMNs) and 500 µg/dl protein. He was discharged after 15 days following empiric treatment with ceftazidime and amikacin. mNGS identified *B. cereus* as the potential etiology. Follow-up at the age of 1 year revealed that the child required ventriculoperitoneal shunt placement for hydrocephalus. The child currently does not have any significant health problems.

### Mumps virus

CHRF0036: A 13-year old boy was admitted after 7 days of fever with irritability and headache. His CSF had 1,500 WBC/µl (60% PMNs) and 160 mg/dl protein. The patient was treated empirically with ceftriaxone for 10 days, after which the family left against medical advice. The logistic regression classifier failed to identify a potential pathogenic microbe. However, manual inspection of the data identified two reads that mapped mumps virus. This was further confirmed with validated qPCR of the original CSF sample to detect Mumps and Sanger sequencing of the resultant amplicon. Mumps virus was not detected in any other samples, except for CHRF0011 (see below). During a follow-up conversation with the father, he reported that the child had had parotitis and fever preceding the headaches and that he had made a full recovery.

CHRF0011: An 18-month old boy was admitted with fever and convulsion. His CSF revealed 100 WBC/ µl (60% PMNs) and 60 µg/dl protein. He was treated with ceftriaxone and discharged after 9 days. mNGS identified mumps virus with sufficient read coverage to determine the genotype as G. At the age of 2.5 years, the child was healthy.

### Enterovirus B

CHRF0010: A 7-month old boy was admitted due to fever, convulsion and lethargy. His CSF analysis unveil 314 WBC/µl (80% PMNs) and 40 µg/dl protein. He was discharged after 6 days of empiric treatment with ceftriaxone. mNGS identified enterovirus B. Almost a year after the episode, the father reported his child falls frequently.

### Chikungunya virus

CHRF0071: A 1-month old girl was admitted with fever, rash, convulsion, diarrhea and lethargy. Her CSF contained 180 WBC/µl (80% PMNs) and 250 µg/dl protein. She was treated with ceftazidime and amikacin and discharged after 6 days. mNGS detected CHIKV with complete genome coverage. The child is currently healthy.

CHRF0094: A 5-day old girl was admitted with fever, convulsion and lethargy. CSF contained 1,000 WBC/µl (90% PMNs) and 220 µg/dl protein. She was treated with ceftazidime and amikacin, followed by meropenem and was discharged after 10 days. mNGS detected CHIKV with complete genome coverage. The child was healthy during our follow up.

CHIRF0012: An 86-month old girl was admitted with acute glomerulonephritis, fever, convulsion, abdominal distension, edema, lethargy and generalized weakness. A LP was performed after 36 days, presumably due to development of meningitis-like symptoms. The CSF contained 180 WBC/µl (60% PMNs) and 55 µg/dl protein. She was treated empirically with cefuroxime, metronidazole, ciprofloxacin, ceftazidime, and acyclovir. The child finally died after 45 days. mNGS identified CHIKV, and the complete genome was assembled.

### Neuroinvasive CHIKV in Bangladesh

All belonging to the East/Central/South African (ECSA) CHIKV genotype, the three CHIKV genomes from the above cases were >99% identical to each other, and to the genome of the strain that caused a febrile outbreak in Dhaka in the summer of 2017. Two of the three children with CHIKV were admitted in June and July of 2017, the peak of the febrile outbreak. To determine if there were additional cases of meningitis admitted in DSH during that period, we performed CHIKV-specific qPCR on 472 idiopathic CSF specimens collected and stored in 2017 and identified 17 additional CHIKV cases. We detected significant overlap between dates of collection of the 20 CHIKV-positive CSF samples with the dates when suspected and confirmed febrile CHIKV cases appeared in our hospitals (Figure 3). Most of these cases originated in Dhaka city, where the febrile outbreak occurred (Figure S2). The median age of these 20 CHIKV-positive patients was 8 months (range: 8 days – 96 months), 35% were female. The mean CSF WBC count was 188/µl (range: 12–1200 WBC/ µl), and the mean PMNs was 48% (range: 10-90%). The average hospital length of stay was 11 days (range: 2-45 days), and the 30 day mortality rate was 0.05% (1/20).

**Figure 3.**
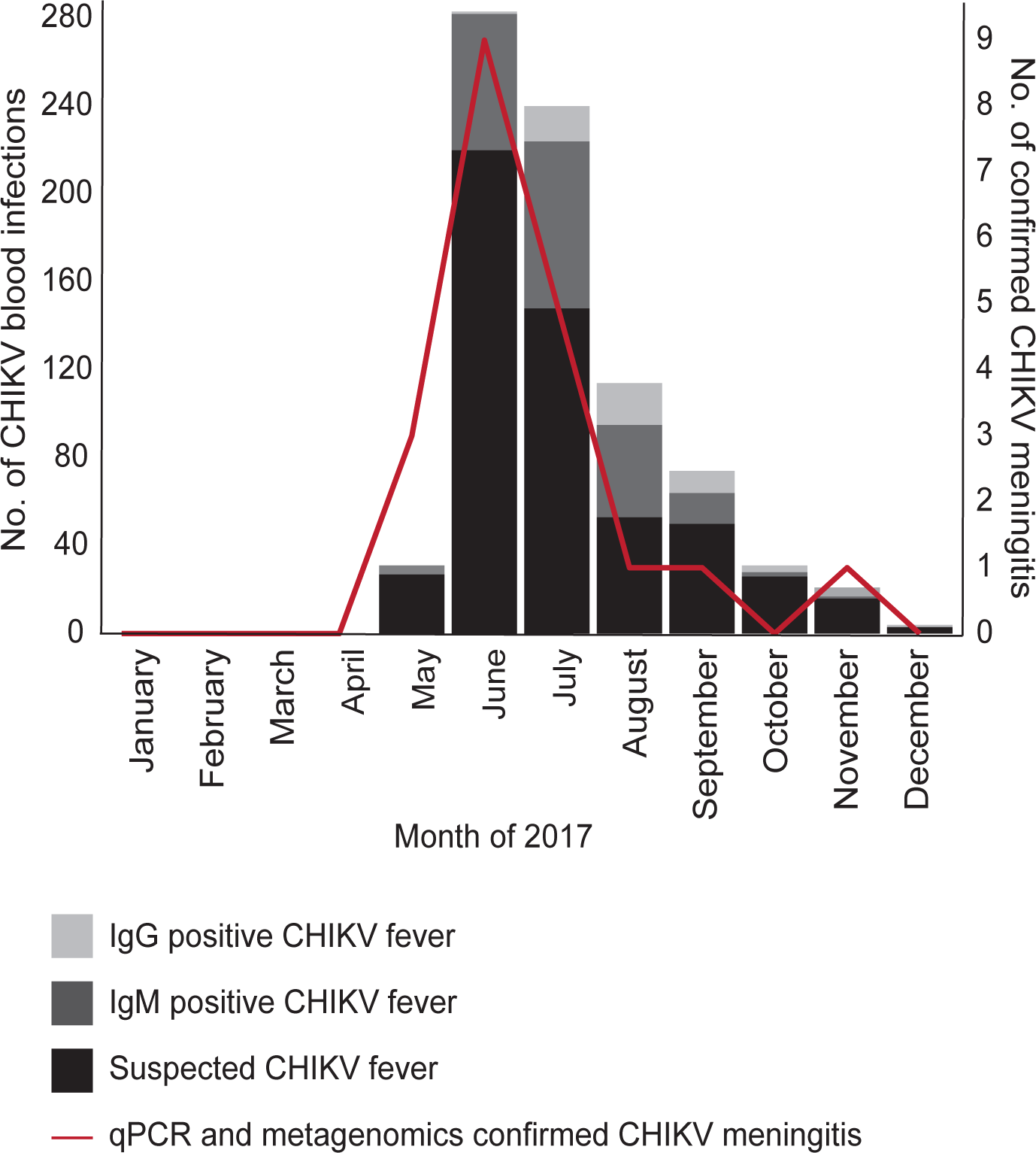
Chikungunya meningitis outbreak in Bangladesh. CHIKV meningitis outbreak overlapped with CHIKV febrile outbreak. The months in 2017 when the CHIKV-positive meningitis CSF samples were collected and when suspected febrile CHIKV cases sought care in the two largest pediatric hospitals of Bangladesh, Dhaka Shishu Hospital and Shishu Shasthya Foundation Hospital. The blood samples of suspected febrile CHIKV cases were detected by a specific diagnostic test for CHIKV-IgG and IgM (SD Biosensor, South Korea) as part of clinical care and results were collected retrospectively from laboratory records.

Subsequent mNGS of CSF from these 17 additional cases identified CHIKV RNA in all (Table S3). Comparison with other CHIKV genomes available in NCBI showed close relationship of the Bangladeshi strain with other strains that caused outbreaks in Asia in recent years, specifically to the one that caused an outbreak in Pakistan in 2016 (99.8% identity (Figure 4).

**Figure 4.**
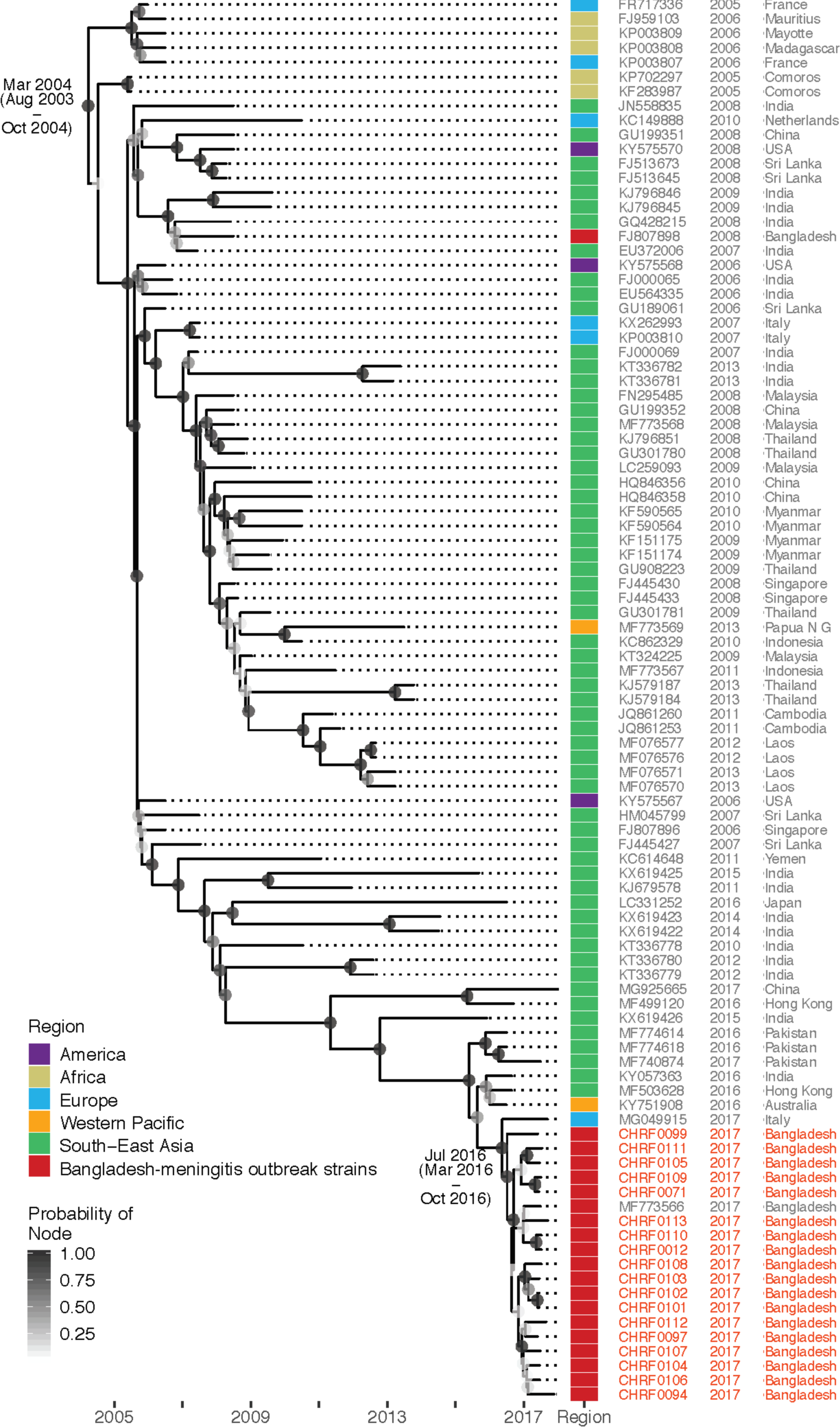
Genetic relationship of Bangladeshi CHIKV-meningitis strain with strains that caused recent outbreak/s in Bangladesh and elsewhere. All CHIV genomes identified and assembled in this study were compared with selected CHIKV genomes available in NCBI data for time-resolved phylogenetics.

## Discussion

Prevention and effective and timely treatment of pediatric meningitis in LMICs is essential for achieving the United Nation’s Sustainable Development Goal 3 of ensuring healthy lives and promoting well-being for all at all ages. Obtaining a microbiological diagnosis can improve outcomes by informing targeted antimicrobial therapy for individual patients. On a population level, improved surveillance better informs region-specific treatment and prevention policy decisions and monitor outbreaks. Here, we coupled unbiased mNGS and machine learning with traditional diagnostics to bridge knowledge gaps and identify the etiology of meningitis in Bangladeshi children.

For samples with known etiology previously determined by qPCR and/or by antigen or serology testing, unbiased mNGS correctly identified the pathogen in 25 of 36 (69.4%) samples. After excluding cases with low pathogen abundance as defined by a confirmatory qPCR Ct of >32, mNGS identified 24 of 27 (89%) known infections. The decreased sensitivity of CSF mNGS for very low abundance pathogens has been previously reported.^22^ A potential pathogen was identified in 40% (10/25) of previously idiopathic meningitis cases. The microbes detected in these cases included bacteria, mycobacteria and viruses with established CNS pathogenicity that were likely not detected due to a variety of reasons including prior antibiotic consumption, the lack of available clinical laboratory assays, and in the case of the CHIKV cases, lack of clinical suspicion for a newly emerging virus with underrecognized neuroinvasive potential.

Among potential bacterial pathogens, we identified *S. enterica* in one sample. Even though that CSF specimen was culture-negative, a culture obtained one week earlier grew *S. enterica*, suggesting that mNGS identified an infection even after antibiotic treatment cleared live pathogens. *B. cereus,* detected in a 6-day old neonate, is commonly recognized as a CNS pathogen in immunocompromised patients including neonates, and at least 7 cases of neonatal *B. cereus* meningitis have been reported.^23,24^ *S. maltophila* was identified in an 4-month old patient; this is a global emerging multidrug resistant gram-negative species of bacteria frequently associated with infections in young or immunocompromised patients^25^. *Mycobacterium tuberculosis*, a well-known cause of severe and chronic meningitis in TB-endemic countries,^26^ was detected from one patient. This finding was strengthened by the patient’s potential exposure in the household and a serendipitous subsequent clinical diagnosis unrelated to this study. The time-consuming nature of culture, the low yield of smear-based detection, makes detection of TB meningitis notoriously challenging and suggests the potential value of mNGS for broad range screening of diverse pathogens including TB.

Among the viral pathogens detected in this study, mumps virus was detected in two cases, an organism well-associated with meningitis.^27,28^ Enterovirus B was detected in one case, also a known cause of pediatric meningitis in the South Asia region.^29^

One of the most interesting findings in this study was the detection of CHIKV in three children, two of which occurred amidst a 2017 CHIKV outbreak in Bangladesh. In recent years, there have been increasing reports of neurological complications associated with CHIKV infection,^30,31^ but no neuroinvasive cases have been identified previously in Bangladesh, and no clinical testing for CHIKV in CSF was being performed in Bangladesh at the time. Our mNGS findings led us to retrospectively screen 472 CSF samples from DSH collected through 2017 with a CHIKV qPCR assay. This identified an additional 17 CHIKV cases, bringing the total to 20. The rate of detection of CHIKV meningitis was consistent with the rate of detection of CHIKV fever outbreak cases reported in Dhaka during the same time period. This study has enhanced the viral detection capabilities at the DSH laboratory, consistent with outbreak preparedness goals. Future studies will include a health-care utilization survey in the catchment area surrounding the DSH to ascertain disease incidence,^32^ in addition to follow-up surveillance for neurodevelopmental assessment of the affected children.

The findings of this study should be considered within the context of several limitations. Importantly, CSF specimens were not collected and stored specifically for metagenomic analysis; this study used samples stored following no stringent or consistent guidelines, with no specific reagents, over a period of six years, which may have affected the quality of the RNA in the samples. Furthermore, this was a retrospective study, and thus, the timing of sample collection with respect to days of disease evolution and prior antibiotic exposure was uncontrolled and unknown in most cases. The quantity of pathogen material may be limiting or undetectable with time, which may contribute to the proportion of unresolved cases in this study. Future studies will include prospective surveillance following improved guidelines for sample collection, storage, processing, and sequencing. The availability of open-source cloud-based pipelines, such as IDseq, for bioinformatic interrogation of metagenomic samples, obviates the need for expensive on-premise compute, storage, and IT personnel support, and future studies can be conducted on site, in real time.

## Conclusions

Hypothesis-free pathogen identification methods like mNGS can facilitate the identification of infectious causes of meningitis that had eluded standard laboratory testing, including previously unrecognized outbreaks of neuroinvasive viral infections like CHIKV. These improved patient and population-level data can inform better health policy decisions, including but not limited to vaccine deployment, antibiotic stewardship, vector control and pandemic preparedness.

## Acknowledgments

We are grateful to Md Shariful Islam, Ms Rhidita Saha, Ms Tasmim Sultana Lipi for assistance with clinical phone follow-up of meningitis cases, to Ms Popy Devanth for assistance with CSF processing and storage in Child Health Research Foundation, to Dr. Mohammad Jamal Uddin, Dr. Shampa Saha and Dr. ASM Nawshad Uddin for clinical and technical support during meningitis surveillance in Dhaka Shishu Hospital. We are also thankful to Maira Phelps and Annie Lo for technical assistance with CSF storage and processing at the Chan Zuckerberg Biohub. Meningitis surveillance in Dhaka Shishu Hospital was supported by Gavi, the Vaccine Alliance, through the World health Organization-supported Invasive Bacterial Vaccine Preventable Diseases study (grant numbers 201588766, 201233523, 201022732, 200749550, 201686542, 202048618 & 202048971). This study was supported by the Bill and Melinda Gates Foundation (OPP1198769).

EDC reports receiving consulting fees and licensing fees from Vela Diagnostics, on topics outside the scope of this work. MRW has received funding from Roche/Genentech on topics outside the scope of this work. SKS has received research grants from GlaxoSmithKline, Sanofi Pasteur, and Pfizer outside the scope of this work. JDR is a board member of uBiome, Inc., and a scientific consultant for Allen & Co., LLC. Other authors have declared no conflict of interest.

Data access, responsibility and analysis: SS and JDR had full access to all the data in the study and takes responsibility for the integrity of the data and the accuracy of the data analysis.

---

**Table 1.** Summary of clinical characteristics of idiopathic cases (n=25).

| Sample ID | Age (months) | WBC/ul (PMN%) / Protein (mg/dl) | Pathogen detected | Provisional/ Final diagnosis | Hospital duration (days)/Outcome | Long-term outcome |
| --- | --- | --- | --- | --- | --- | --- |
| **Resolved** |  |  |  |  |  |  |
| CHRF0010 | 7/m | 314 (80) / 40 | Enterovirus B | Meningitis/Meningitis | 6/Dis | Heathy |
| CHRF0058 | 160/f | 360 (80) / 200 | *M. tuberculosis* | ICSOL/ ICSOL | 15/LAMA | Sequalae |
| CHRF0070 | 0/m | 12000 (95) / 500 | *B. cereus* | Sepsis | 15/Dis | Heathy |
| CHRF0082 | 4/m | 90 (70) / 600 | *S. enterica* | Meningitis/Meningitis | 23/Dis | Healthy |
| CHRF0094 | 0/f | 1000 (90) / 220 | CHIKV | Meningitis/Meningitis | 10/Dis | Healthy |
| CHRF0011 | 18/m | 100 (60) / 60 | Mumps | Meningitis/Meningitis | 9/Dis | Healthy |
| CHRF0035 | 21/f | 600 (90) / 700 | Human herpes virus 6 | Acute stroke syndrome/ Acute stroke syndrome | 56/Died | Died |
| CHRF0059 | 4/f | 120 (80) / 300 | *S. maltophilia* | Meningitis/Meningitis | 28/Dis | NA |
| CHRF0071 | 1/f | 180 (80) / 250 | CHIKV | Meningitis/Meningitis | 6/Dis | Heathy |
| CHRF0012 | 86/f | 180 (60) / 55 | CHIKV | Acute Glomerulonephritis/Meningoencephalitis | 45/Died | Died |
| CHRF0036 | 156/m | 1500 (60) / 160 | Mumps | Meningitis/Meningitis | 10/LAMA | Healthy |
| **Unresolved** |  |  |  |  |  |  |
| CHRF0081 | 72/m | 1100 (40) / 350 | NA | Meningitis/Meningitis | 37/LAMA | Died |
| CHRF0093 | 13/m | 20 (80) / 80 | NA | Heart disease/ Dextocardia | 11/Died | Died |
| CHRF0022 | 3/m | 460 (80) / 300 | NA | Meningitis/Meningitis | 9/LAMA | Died |
| CHRF0034 | 23/f | 70 (70) / 150 | NA | Meningitis/Meningitis | 11/Dis | Sequelae |
| CHRF0046 | 1/f | 160 (80) / 70 | NA | Pneumonia/ Pneumonia | 4/LAMA | Healthy |
| CHRF0023 | 4/m | 74 (70) / 100 | NA | Meningitis/ Meningitis | 8/Dis | Sequelae |
| CHRF0047 | 16/m | 2600 (90) / 400 | NA | ARI/Meningoencephalitis | 5/Died | Died |
| CHRF0083 | 7/m | 5600 (80) / 1000 | NA | Hydrocephalus/Pneumonia | 10/Died | Died |
| CHRF0095 | 98/m | 90 (70) / 60 | NA | Meningoencephalitis/Tubercular meningitis | 33/LAMA | Died |
| CHRF0024 | 96/f | 460 (60) / 200 | NA | Meningitis/Meningitis | 8/LAMA | NA |
| CHRF0048 | 2/m | 3800 (80) / 400 | NA | Meningitis/Meningitis | 35/Dis | NA |
| CHRF0060 | 4/m | 84 (70) / 150 | NA | Meningitis/Meningitis | 11/LAMA | Died |
| CHRF0072 | 3/f | 900 (60) / 300 | NA | Meningitis/Meningitis | 44/Dis | Sequelae |
| CHRF0084 | 4/f | 70 (60) / 250 | NA | Meningitis/Meningitis | 51/Died | Died |

NA: not available; LAMA: Left against medical advice; Dis: Discharged; ICSOL: intracranial space-occupying lesion. NA: not available

## Supplemental Material

Method S1. **Bioinformatic analysis and pathogen identification.**

Method S2. **Phylogenetic analysis of Chikungunya virus strains responsible for the meningitis outbreak in Bangladesh, 2017.**

**Figure S1.**
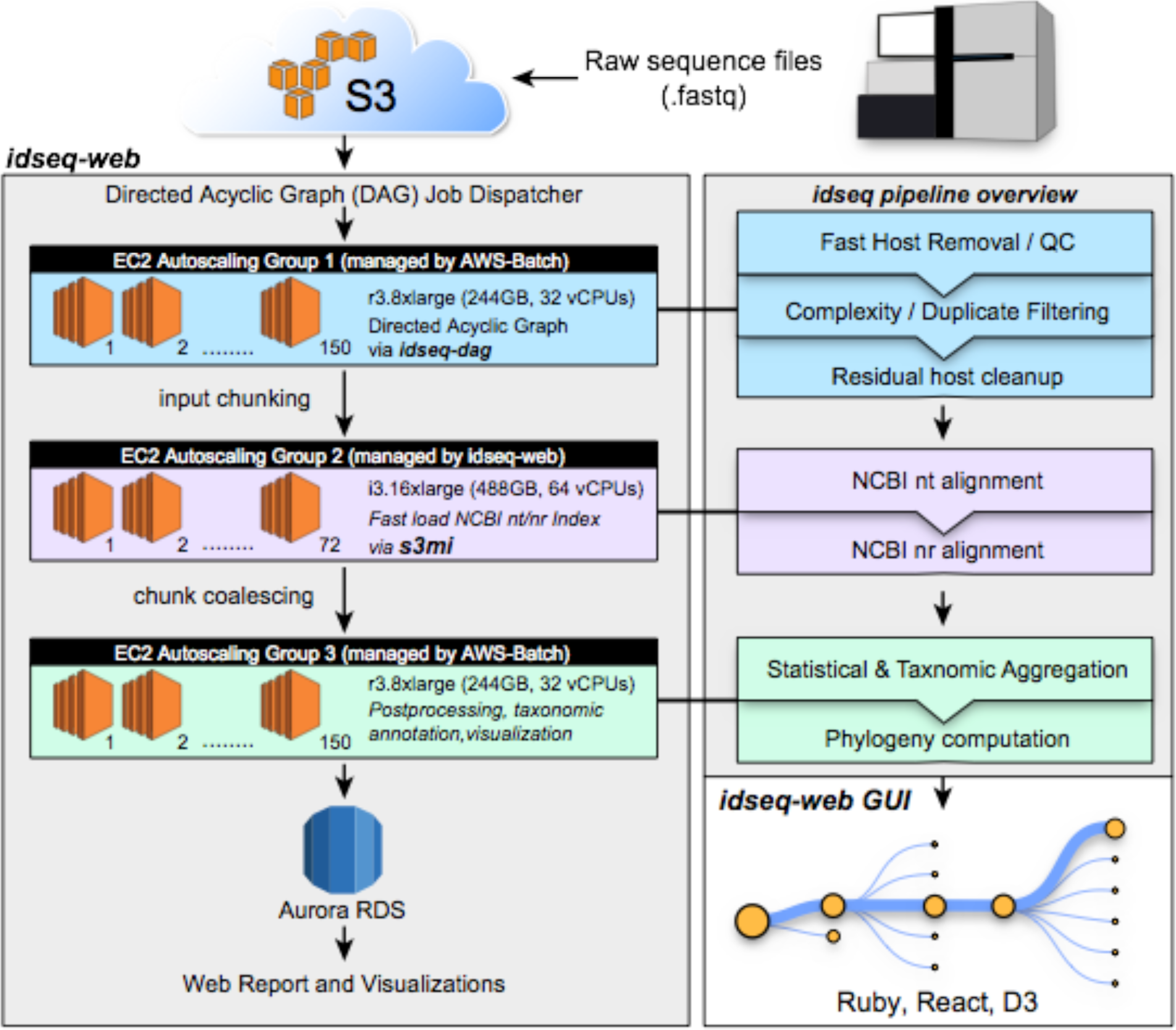
Schematic representation of the bioinformatic approach in IDSEQ for pathogen identification.

**Figure S2.**
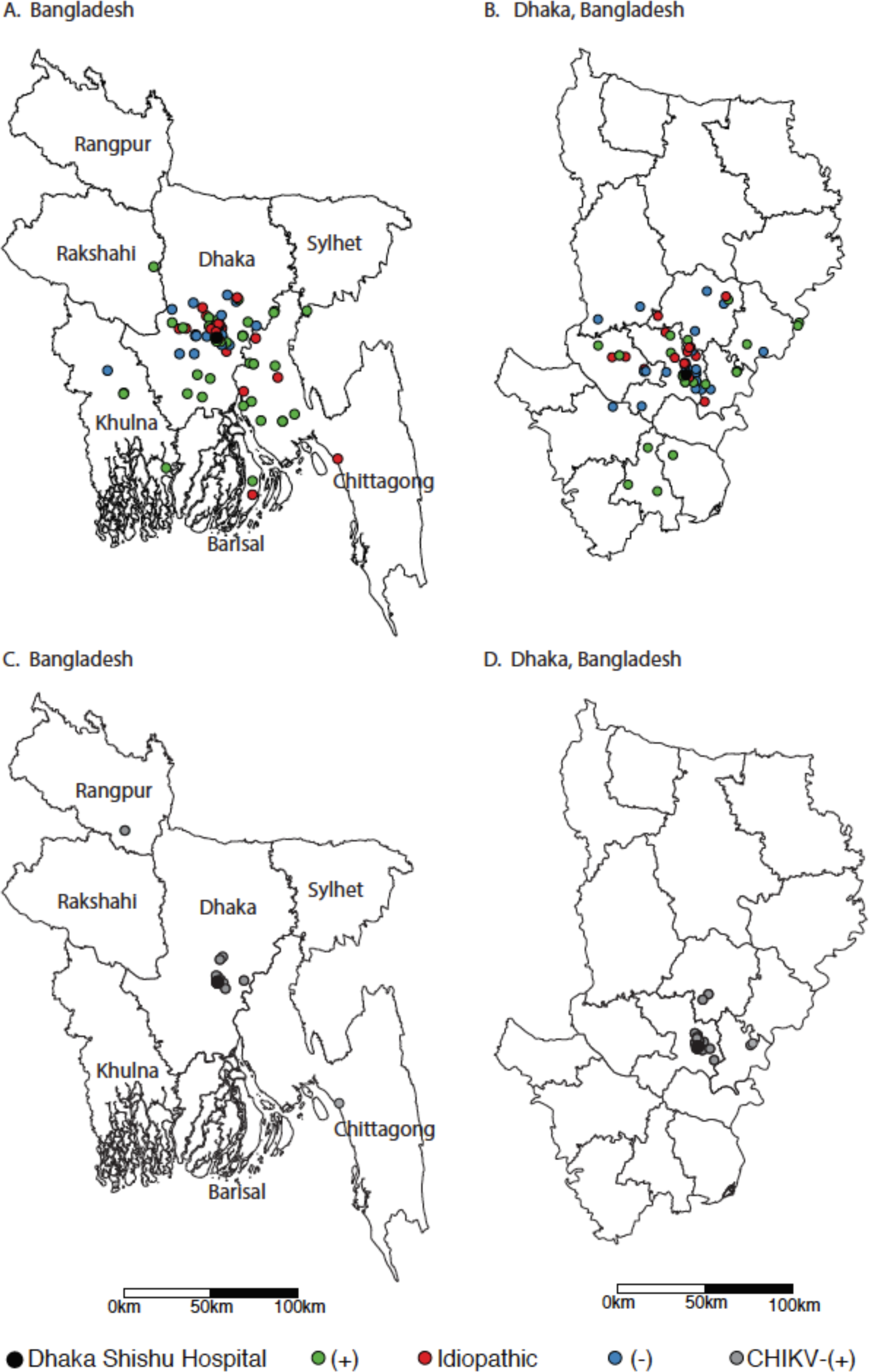
Residential locations of all selected cases and CHIKV-positive meningitis cases. A. The map of Bangladesh depicting national distribution of all cases. B. Magnified map of Dhaka division from where majority of cases arrived. C The map of Bangladesh depicting national distribution of all CHIKV-positive meningitis cases during the outbreak of 2017. D. Magnified map of Dhaka division from where almost of CHIKV-meningitis cases arrived from during the outbreak in 2017.

**Table S1.** WHO-defined clinical criteria used to enroll children in the meningitis surveillance in Dhaka Shishu Hospital, Bangladesh.

| WHO case definition | Inclusion criteria |
| --- | --- |
| Meningitis | Any child aged 0-59 months hospitalized with sudden onset of fever >100.4 °F and one of the following signs:<br>• Stiff neck<br>• Altered or reduced level of consciousness<br>• Bulging fontanel (if < 12 months of age)<br>• Prostration/lethargy<br>• Convulsions<br>• Toxic appearance<br>• Petechial or purpuric rash<br>• Poor Sucking<br>• Irritability (>2 months)<br>Or any child aged 0-59 months hospitalized with a clinical diagnosis of meningitis. |

**Table S2.**
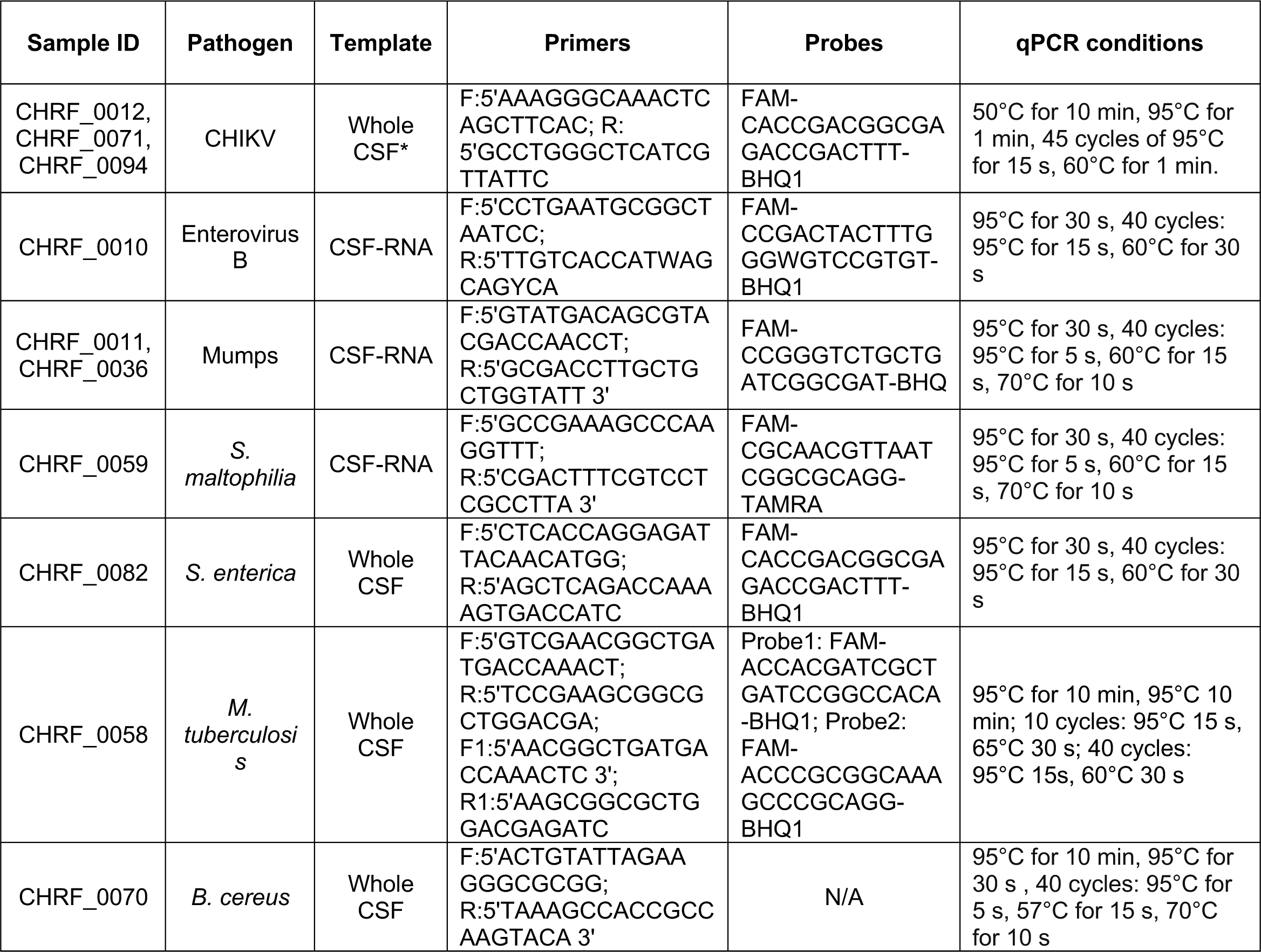
Detection of Chikungunya virus and orthogonal confirmation of probable pathogens through qPCR. For CHIKV confirmation, we used qScript XLT 1-Step RT-qPCR ToughMix Low-ROX (Cat. No. 95134-100). For the remaining confirmations, we used Quantabio PerfeCTa qPCR ToughMix, Low Rox (Cat. No. 95114-012). For reverse transcription, we used Quantabio qScript cDNA Super Mix (Cat. No. 95048-100): 25°C for 5 min, 42°C for 30 min, 85°C for 5 min.

**Table S3.**
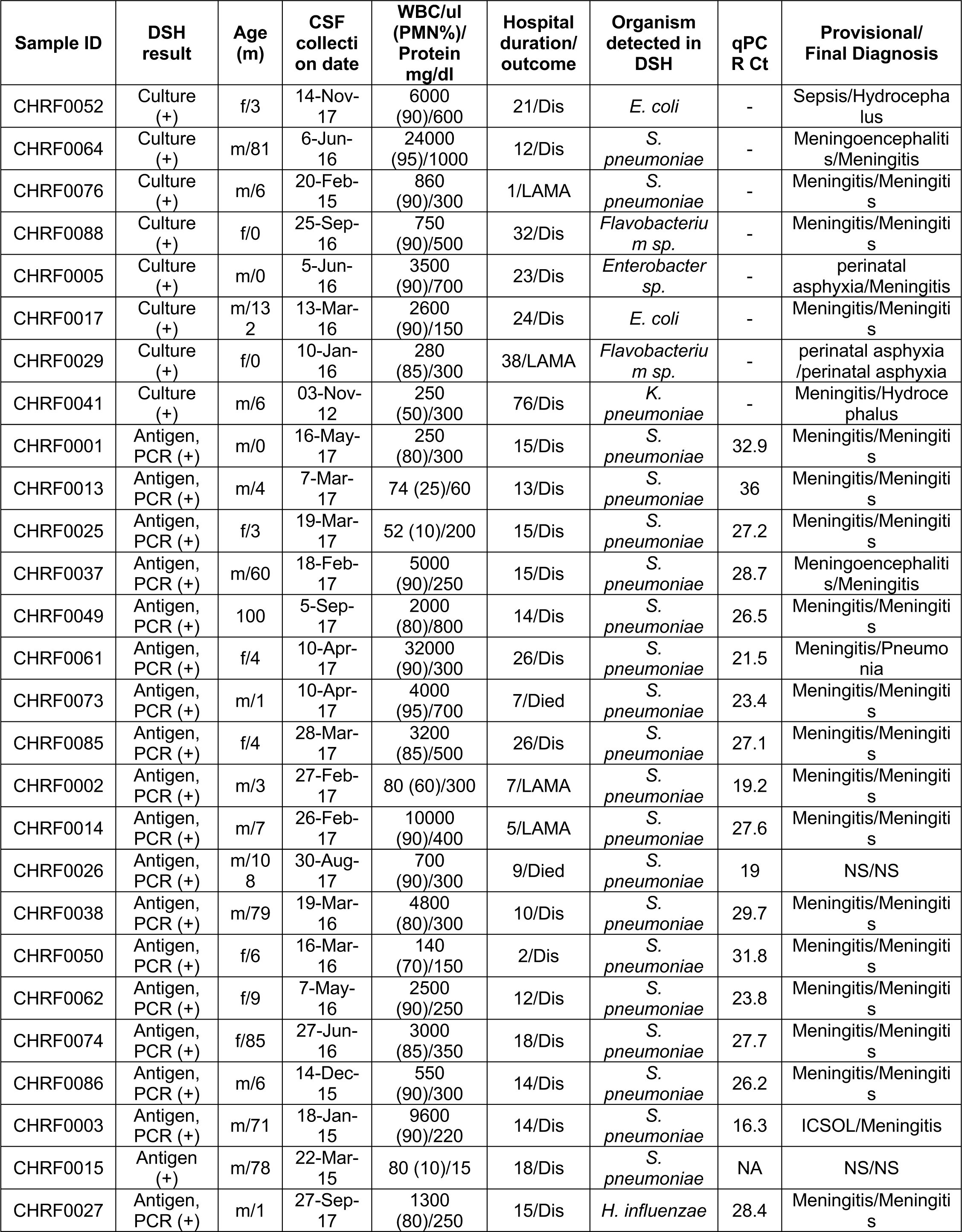
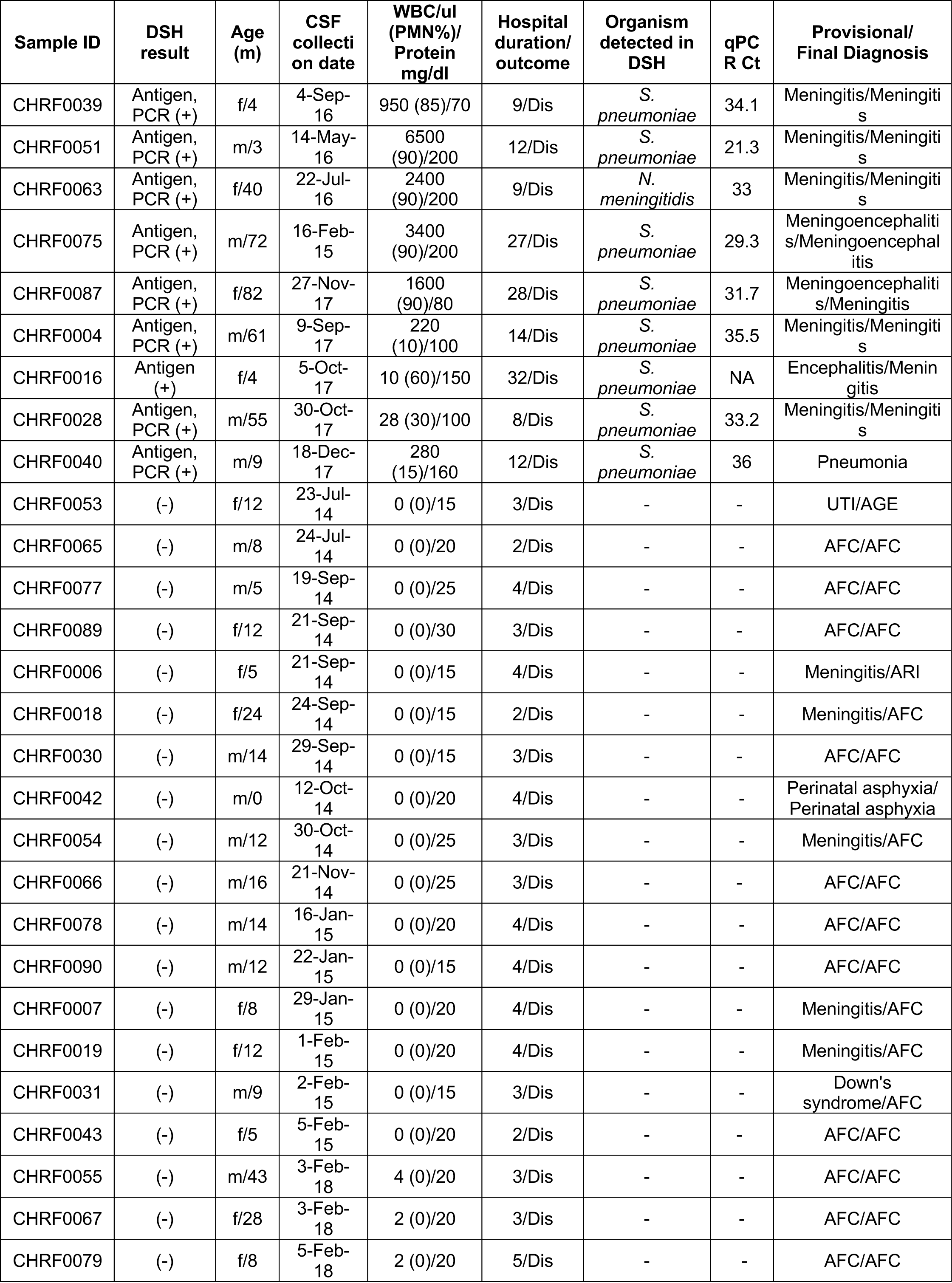
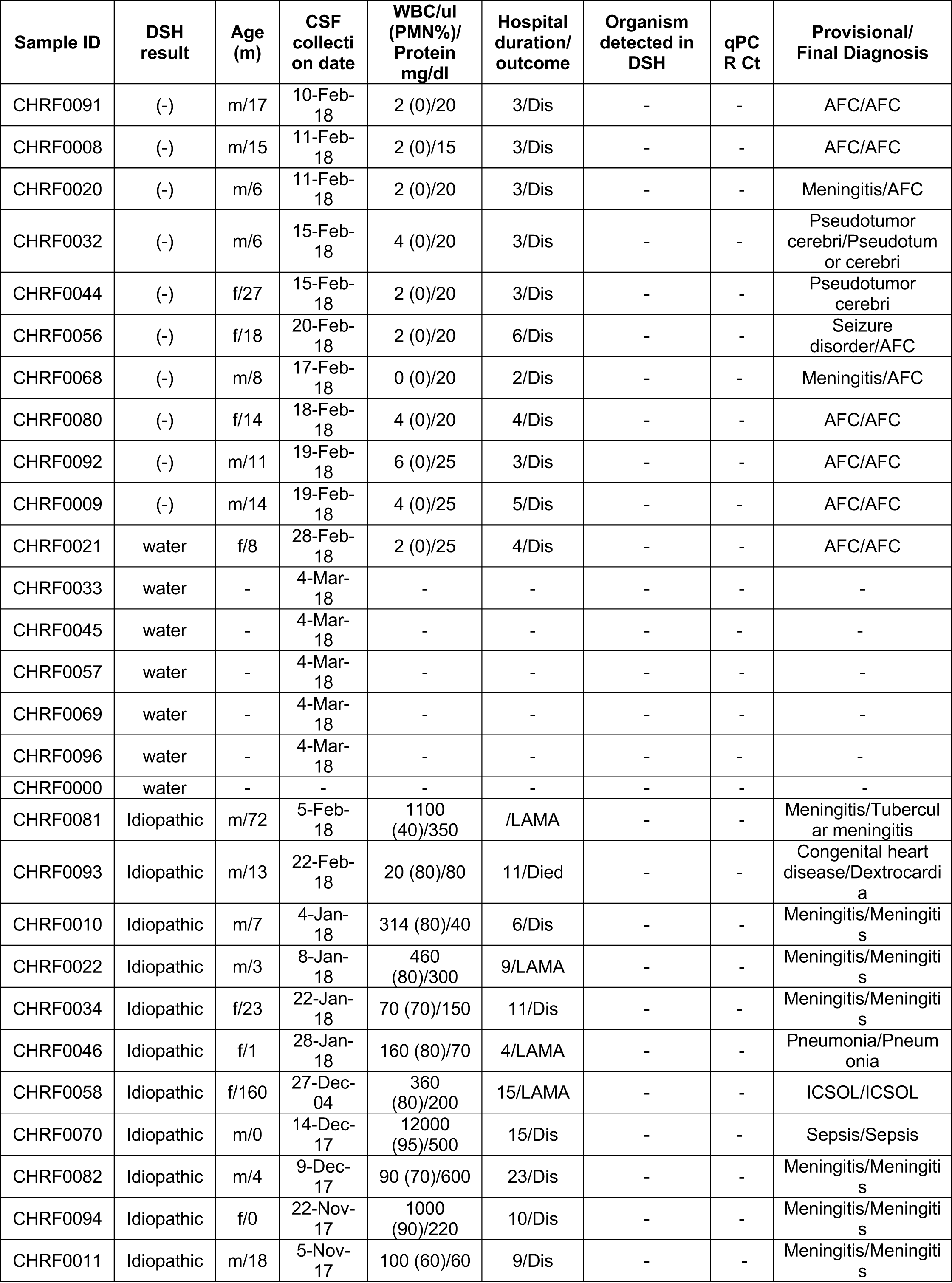
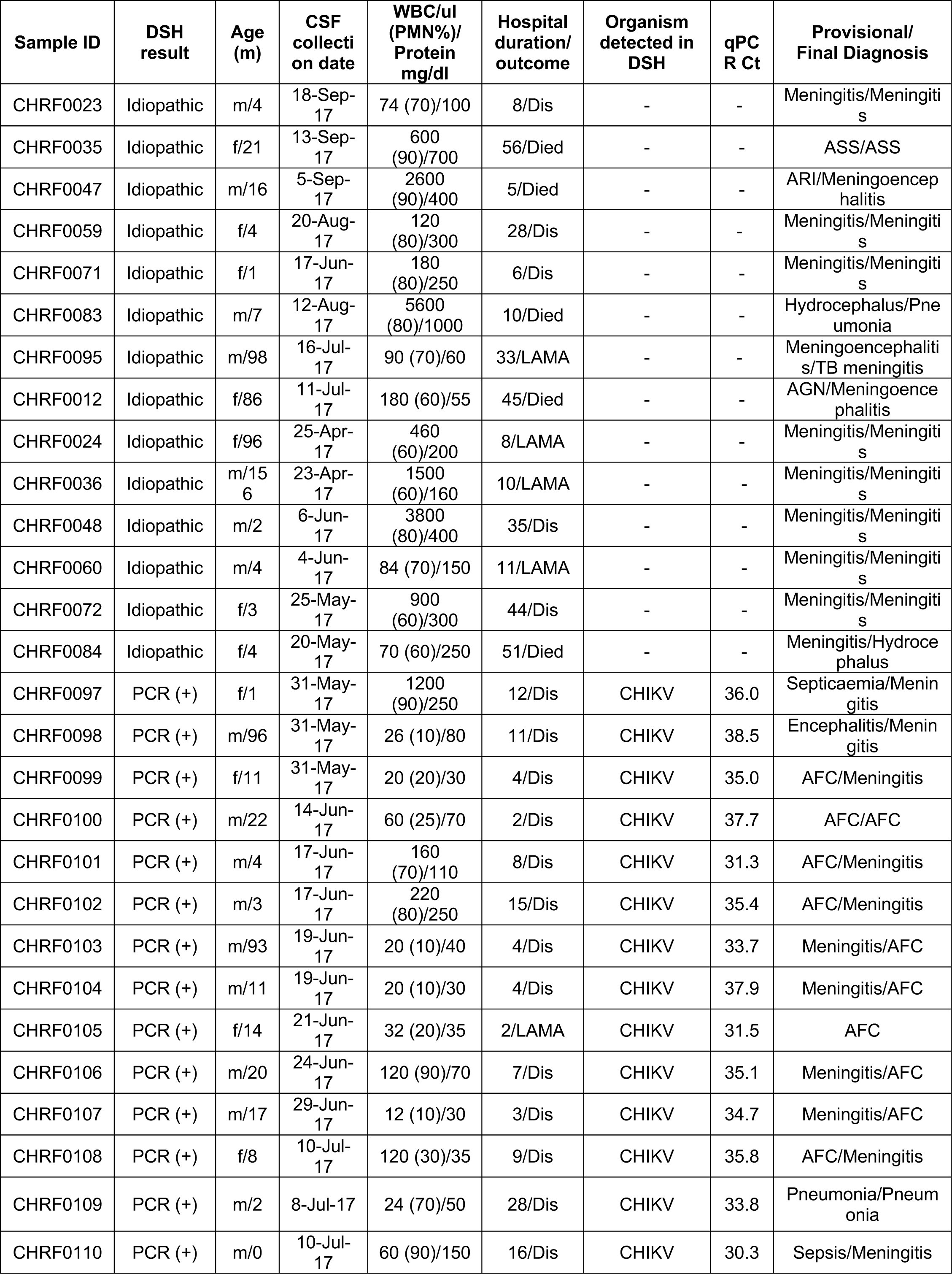
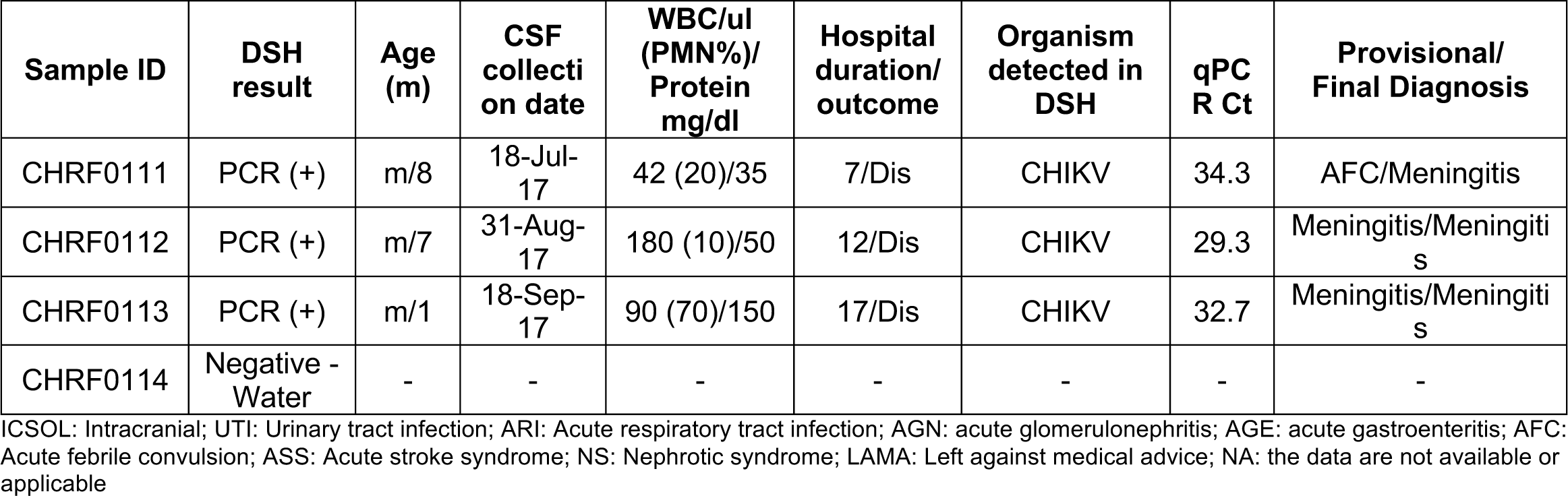
Case-based clinical and laboratory metadata of all cases included in this study (n=115).

| Sample ID | DSH result | Age (m) | CSF collecti on date | WBC/ul (PMN%)/ Protein mg/dl | Hospital duration/ outcome | Organism detected in DSH | qPC R Ct | Provisional/ Final Diagnosis |
| --- | --- | --- | --- | --- | --- | --- | --- | --- |
| CHRF0052 | Culture (+) | f/3 | 14-Nov-17 | 6000 (90)/600 | 21/Dis | *E. coli* | - | Sepsis/Hydrocephalus |
| CHRF0064 | Culture (+) | m/81 | 6-Jun-16 | 24000 (95)/1000 | 12/Dis | *S. pneumoniae* | - | Meningoencephalitis/Meningitis |
| CHRF0076 | Culture (+) | m/6 | 20-Feb-15 | 860 (90)/300 | 1/LAMA | *S. pneumoniae* | - | Meningitis/Meningitis |
| CHRF0088 | Culture (+) | f/0 | 25-Sep-16 | 750 (90)/500 | 32/Dis | *Flavobacterium* sp. | - | Meningitis/Meningitis |
| CHRF0005 | Culture (+) | m/0 | 5-Jun-16 | 3500 (90)/700 | 23/Dis | *Enterobacter* sp. | - | perinatal asphyxia/Meningitis |
| CHRF0017 | Culture (+) | m/132 | 13-Mar-16 | 2600 (90)/150 | 24/Dis | *E. coli* | - | Meningitis/Meningitis |
| CHRF0029 | Culture (+) | f/0 | 10-Jan-16 | 280 (85)/300 | 38/LAMA | *Flavobacterium* sp. | - | perinatal asphyxia /perinatal asphyxia |
| CHRF0041 | Culture (+) | m/6 | 03-Nov-12 | 250 (50)/300 | 76/Dis | *K. pneumoniae* | - | Meningitis/Hydrocephalus |
| CHRF0001 | Antigen, PCR (+) | m/0 | 16-May-17 | 250 (80)/300 | 15/Dis | *S. pneumoniae* | 32.9 | Meningitis/Meningitis |
| CHRF0013 | Antigen, PCR (+) | m/4 | 7-Mar-17 | 74 (25)/60 | 13/Dis | *S. pneumoniae* | 36 | Meningitis/Meningitis |
| CHRF0025 | Antigen, PCR (+) | f/3 | 19-Mar-17 | 52 (10)/200 | 15/Dis | *S. pneumoniae* | 27.2 | Meningitis/Meningitis |
| CHRF0037 | Antigen, PCR (+) | m/60 | 18-Feb-17 | 5000 (90)/250 | 15/Dis | *S. pneumoniae* | 28.7 | Meningoencephalitis/Meningitis |
| CHRF0049 | Antigen, PCR (+) | 100 | 5-Sep-17 | 2000 (80)/800 | 14/Dis | *S. pneumoniae* | 26.5 | Meningitis/Meningitis |
| CHRF0061 | Antigen, PCR (+) | f/4 | 10-Apr-17 | 32000 (90)/300 | 26/Dis | *S. pneumoniae* | 21.5 | Meningitis/Pneumonia |
| CHRF0073 | Antigen, PCR (+) | m/1 | 10-Apr-17 | 4000 (95)/700 | 7/Died | *S. pneumoniae* | 23.4 | Meningitis/Meningitis |
| CHRF0085 | Antigen, PCR (+) | f/4 | 28-Mar-17 | 3200 (85)/500 | 26/Dis | *S. pneumoniae* | 27.1 | Meningitis/Meningitis |
| CHRF0002 | Antigen, PCR (+) | m/3 | 27-Feb-17 | 80 (60)/300 | 7/LAMA | *S. pneumoniae* | 19.2 | Meningitis/Meningitis |
| CHRF0014 | Antigen, PCR (+) | m/7 | 26-Feb-17 | 10000 (90)/400 | 5/LAMA | *S. pneumoniae* | 27.6 | Meningitis/Meningitis |
| CHRF0026 | Antigen, PCR (+) | m/108 | 30-Aug-17 | 700 (90)/300 | 9/Died | *S. pneumoniae* | 19 | NS/NS |
| CHRF0038 | Antigen, PCR (+) | m/79 | 19-Mar-16 | 4800 (80)/300 | 10/Dis | *S. pneumoniae* | 29.7 | Meningitis/Meningitis |
| CHRF0050 | Antigen, PCR (+) | f/6 | 16-Mar-16 | 140 (70)/150 | 2/Dis | *S. pneumoniae* | 31.8 | Meningitis/Meningitis |
| CHRF0062 | Antigen, PCR (+) | f/9 | 7-May-16 | 2500 (90)/250 | 12/Dis | *S. pneumoniae* | 23.8 | Meningitis/Meningitis |
| CHRF0074 | Antigen, PCR (+) | f/85 | 27-Jun-16 | 3000 (85)/350 | 18/Dis | *S. pneumoniae* | 27.7 | Meningitis/Meningitis |
| CHRF0086 | Antigen, PCR (+) | m/6 | 14-Dec-15 | 550 (90)/300 | 14/Dis | *S. pneumoniae* | 26.2 | Meningitis/Meningitis |
| CHRF0003 | Antigen, PCR (+) | m/71 | 18-Jan-15 | 9600 (90)/220 | 14/Dis | *S. pneumoniae* | 16.3 | ICSOL/Meningitis |
| CHRF0015 | Antigen (+) | m/78 | 22-Mar-15 | 80 (10)/15 | 18/Dis | *S. pneumoniae* | NA | NS/NS |
| CHRF0027 | Antigen, PCR (+) | m/1 | 27-Sep-17 | 1300 (80)/250 | 15/Dis | *H. influenzae* | 28.4 | Meningitis/Meningitis |
| CHRF0039 | Antigen, PCR (+) | f/4 | 4-Sep-16 | 950 (85)/70 | 9/Dis | *S. pneumoniae* | 34.1 | Meningitis/Meningiti s |
| CHRF0051 | Antigen, PCR (+) | m/3 | 14-May-16 | 6500 (90)/200 | 12/Dis | *S. pneumoniae* | 21.3 | Meningitis/Meningiti s |
| CHRF0063 | Antigen, PCR (+) | f/40 | 22-Jul-16 | 2400 (90)/200 | 9/Dis | *N. meningitidis* | 33 | Meningitis/Meningiti s |
| CHRF0075 | Antigen, PCR (+) | m/72 | 16-Feb-15 | 3400 (90)/200 | 27/Dis | *S. pneumoniae* | 29.3 | Meningoencephaliti s/Meningoencephal itis |
| CHRF0087 | Antigen, PCR (+) | f/82 | 27-Nov-17 | 1600 (90)/80 | 28/Dis | *S. pneumoniae* | 31.7 | Meningoencephaliti s/Meningitis |
| CHRF0004 | Antigen, PCR (+) | m/61 | 9-Sep-17 | 220 (10)/100 | 14/Dis | *S. pneumoniae* | 35.5 | Meningitis/Meningiti s |
| CHRF0016 | Antigen (+) | f/4 | 5-Oct-17 | 10 (60)/150 | 32/Dis | *S. pneumoniae* | NA | Encephalitis/Menin gitis |
| CHRF0028 | Antigen, PCR (+) | m/55 | 30-Oct-17 | 28 (30)/100 | 8/Dis | *S. pneumoniae* | 33.2 | Meningitis/Meningiti s |
| CHRF0040 | Antigen, PCR (+) | m/9 | 18-Dec-17 | 280 (15)/160 | 12/Dis | *S. pneumoniae* | 36 | Pneumonia |
| CHRF0053 | (-) | f/12 | 23-Jul-14 | 0 (0)/15 | 3/Dis | - | - | UTI/AGE |
| CHRF0065 | (-) | m/8 | 24-Jul-14 | 0 (0)/20 | 2/Dis | - | - | AFC/AFC |
| CHRF0077 | (-) | m/5 | 19-Sep-14 | 0 (0)/25 | 4/Dis | - | - | AFC/AFC |
| CHRF0089 | (-) | f/12 | 21-Sep-14 | 0 (0)/30 | 3/Dis | - | - | AFC/AFC |
| CHRF0006 | (-) | f/5 | 21-Sep-14 | 0 (0)/15 | 4/Dis | - | - | Meningitis/ARI |
| CHRF0018 | (-) | f/24 | 24-Sep-14 | 0 (0)/15 | 2/Dis | - | - | Meningitis/AFC |
| CHRF0030 | (-) | m/14 | 29-Sep-14 | 0 (0)/15 | 3/Dis | - | - | AFC/AFC |
| CHRF0042 | (-) | m/0 | 12-Oct-14 | 0 (0)/20 | 4/Dis | - | - | Perinatal asphyxia/ Perinatal asphyxia |
| CHRF0054 | (-) | m/12 | 30-Oct-14 | 0 (0)/25 | 3/Dis | - | - | Meningitis/AFC |
| CHRF0066 | (-) | m/16 | 21-Nov-14 | 0 (0)/25 | 3/Dis | - | - | AFC/AFC |
| CHRF0078 | (-) | m/14 | 16-Jan-15 | 0 (0)/20 | 4/Dis | - | - | AFC/AFC |
| CHRF0090 | (-) | m/12 | 22-Jan-15 | 0 (0)/15 | 4/Dis | - | - | AFC/AFC |
| CHRF0007 | (-) | f/8 | 29-Jan-15 | 0 (0)/20 | 4/Dis | - | - | Meningitis/AFC |
| CHRF0019 | (-) | f/12 | 1-Feb-15 | 0 (0)/20 | 4/Dis | - | - | Meningitis/AFC |
| CHRF0031 | (-) | m/9 | 2-Feb-15 | 0 (0)/15 | 3/Dis | - | - | Down's syndrome/AFC |
| CHRF0043 | (-) | f/5 | 5-Feb-15 | 0 (0)/20 | 2/Dis | - | - | AFC/AFC |
| CHRF0055 | (-) | m/43 | 3-Feb-18 | 4 (0)/20 | 3/Dis | - | - | AFC/AFC |
| CHRF0067 | (-) | f/28 | 3-Feb-18 | 2 (0)/20 | 3/Dis | - | - | AFC/AFC |
| CHRF0079 | (-) | f/8 | 5-Feb-18 | 2 (0)/20 | 5/Dis | - | - | AFC/AFC |
| CHRF0091 | (-) | m/17 | 10-Feb-18 | 2 (0)/20 | 3/Dis | - | - | AFC/AFC |
| CHRF0008 | (-) | m/15 | 11-Feb-18 | 2 (0)/15 | 3/Dis | - | - | AFC/AFC |
| CHRF0020 | (-) | m/6 | 11-Feb-18 | 2 (0)/20 | 3/Dis | - | - | Meningitis/AFC |
| CHRF0032 | (-) | m/6 | 15-Feb-18 | 4 (0)/20 | 3/Dis | - | - | Pseudotumor cerebri/Pseudotum or cerebri |
| CHRF0044 | (-) | f/27 | 15-Feb-18 | 2 (0)/20 | 3/Dis | - | - | Pseudotumor cerebri |
| CHRF0056 | (-) | f/18 | 20-Feb-18 | 2 (0)/20 | 6/Dis | - | - | Seizure disorder/AFC |
| CHRF0068 | (-) | m/8 | 17-Feb-18 | 0 (0)/20 | 2/Dis | - | - | Meningitis/AFC |
| CHRF0080 | (-) | f/14 | 18-Feb-18 | 4 (0)/20 | 4/Dis | - | - | AFC/AFC |
| CHRF0092 | (-) | m/11 | 19-Feb-18 | 6 (0)/25 | 3/Dis | - | - | AFC/AFC |
| CHRF0009 | (-) | m/14 | 19-Feb-18 | 4 (0)/25 | 5/Dis | - | - | AFC/AFC |
| CHRF0021 | water | f/8 | 28-Feb-18 | 2 (0)/25 | 4/Dis | - | - | AFC/AFC |
| CHRF0033 | water | - | 4-Mar-18 | - | - | - | - | - |
| CHRF0045 | water | - | 4-Mar-18 | - | - | - | - | - |
| CHRF0057 | water | - | 4-Mar-18 | - | - | - | - | - |
| CHRF0069 | water | - | 4-Mar-18 | - | - | - | - | - |
| CHRF0096 | water | - | 4-Mar-18 | - | - | - | - | - |
| CHRF0000 | water | - | - | - | - | - | - | - |
| CHRF0081 | Idiopathic | m/72 | 5-Feb-18 | 1100 (40)/350 | /LAMA | - | - | Meningitis/Tubercul ar meningitis |
| CHRF0093 | Idiopathic | m/13 | 22-Feb-18 | 20 (80)/80 | 11/Died | - | - | Congenital heart disease/Dextrocardi a |
| CHRF0010 | Idiopathic | m/7 | 4-Jan-18 | 314 (80)/40 | 6/Dis | - | - | Meningitis/Meningiti s |
| CHRF0022 | Idiopathic | m/3 | 8-Jan-18 | 460 (80)/300 | 9/LAMA | - | - | Meningitis/Meningiti s |
| CHRF0034 | Idiopathic | f/23 | 22-Jan-18 | 70 (70)/150 | 11/Dis | - | - | Meningitis/Meningiti s |
| CHRF0046 | Idiopathic | f/1 | 28-Jan-18 | 160 (80)/70 | 4/LAMA | - | - | Pneumonia/Pneum onia |
| CHRF0058 | Idiopathic | f/160 | 27-Dec-04 | 360 (80)/200 | 15/LAMA | - | - | ICSOL/ICSOL |
| CHRF0070 | Idiopathic | m/0 | 14-Dec-17 | 12000 (95)/500 | 15/Dis | - | - | Sepsis/Sepsis |
| CHRF0082 | Idiopathic | m/4 | 9-Dec-17 | 90 (70)/600 | 23/Dis | - | - | Meningitis/Meningiti s |
| CHRF0094 | Idiopathic | f/0 | 22-Nov-17 | 1000 (90)/220 | 10/Dis | - | - | Meningitis/Meningiti s |
| CHRF0011 | Idiopathic | m/18 | 5-Nov-17 | 100 (60)/60 | 9/Dis | - | - | Meningitis/Meningiti s |
| CHRF0023 | Idiopathic | m/4 | 18-Sep-17 | 74 (70)/100 | 8/Dis | - | - | Meningitis/Meningiti s |
| CHRF0035 | Idiopathic | f/21 | 13-Sep-17 | 600 (90)/700 | 56/Died | - | - | ASS/ASS |
| CHRF0047 | Idiopathic | m/16 | 5-Sep-17 | 2600 (90)/400 | 5/Died | - | - | ARI/Meningoencep halitis |
| CHRF0059 | Idiopathic | f/4 | 20-Aug-17 | 120 (80)/300 | 28/Dis | - | - | Meningitis/Meningiti s |
| CHRF0071 | Idiopathic | f/1 | 17-Jun-17 | 180 (80)/250 | 6/Dis | - | - | Meningitis/Meningiti s |
| CHRF0083 | Idiopathic | m/7 | 12-Aug-17 | 5600 (80)/1000 | 10/Died | - | - | Hydrocephalus/Pne umonia |
| CHRF0095 | Idiopathic | m/98 | 16-Jul-17 | 90 (70)/60 | 33/LAMA | - | - | Meningoencephaliti s/TB meningitis |
| CHRF0012 | Idiopathic | f/86 | 11-Jul-17 | 180 (60)/55 | 45/Died | - | - | AGN/Meningoence phalitis |
| CHRF0024 | Idiopathic | f/96 | 25-Apr-17 | 460 (60)/200 | 8/LAMA | - | - | Meningitis/Meningiti s |
| CHRF0036 | Idiopathic | m/15 6 | 23-Apr-17 | 1500 (60)/160 | 10/LAMA | - | - | Meningitis/Meningiti s |
| CHRF0048 | Idiopathic | m/2 | 6-Jun-17 | 3800 (80)/400 | 35/Dis | - | - | Meningitis/Meningiti s |
| CHRF0060 | Idiopathic | m/4 | 4-Jun-17 | 84 (70)/150 | 11/LAMA | - | - | Meningitis/Meningiti s |
| CHRF0072 | Idiopathic | f/3 | 25-May-17 | 900 (60)/300 | 44/Dis | - | - | Meningitis/Meningiti s |
| CHRF0084 | Idiopathic | f/4 | 20-May-17 | 70 (60)/250 | 51/Died | - | - | Meningitis/Hydroce phalus |
| CHRF0097 | PCR (+) | f/1 | 31-May-17 | 1200 (90)/250 | 12/Dis | CHIKV | 36.0 | Septicaemia/Menin gitis |
| CHRF0098 | PCR (+) | m/96 | 31-May-17 | 26 (10)/80 | 11/Dis | CHIKV | 38.5 | Encephalitis/Menin gitis |
| CHRF0099 | PCR (+) | f/11 | 31-May-17 | 20 (20)/30 | 4/Dis | CHIKV | 35.0 | AFC/Meningitis |
| CHRF0100 | PCR (+) | m/22 | 14-Jun-17 | 60 (25)/70 | 2/Dis | CHIKV | 37.7 | AFC/AFC |
| CHRF0101 | PCR (+) | m/4 | 17-Jun-17 | 160 (70)/110 | 8/Dis | CHIKV | 31.3 | AFC/Meningitis |
| CHRF0102 | PCR (+) | m/3 | 17-Jun-17 | 220 (80)/250 | 15/Dis | CHIKV | 35.4 | AFC/Meningitis |
| CHRF0103 | PCR (+) | m/93 | 19-Jun-17 | 20 (10)/40 | 4/Dis | CHIKV | 33.7 | Meningitis/AFC |
| CHRF0104 | PCR (+) | m/11 | 19-Jun-17 | 20 (10)/30 | 4/Dis | CHIKV | 37.9 | Meningitis/AFC |
| CHRF0105 | PCR (+) | f/14 | 21-Jun-17 | 32 (20)/35 | 2/LAMA | CHIKV | 31.5 | AFC |
| CHRF0106 | PCR (+) | m/20 | 24-Jun-17 | 120 (90)/70 | 7/Dis | CHIKV | 35.1 | Meningitis/AFC |
| CHRF0107 | PCR (+) | m/17 | 29-Jun-17 | 12 (10)/30 | 3/Dis | CHIKV | 34.7 | Meningitis/AFC |
| CHRF0108 | PCR (+) | f/8 | 10-Jul-17 | 120 (30)/35 | 9/Dis | CHIKV | 35.8 | AFC/Meningitis |
| CHRF0109 | PCR (+) | m/2 | 8-Jul-17 | 24 (70)/50 | 28/Dis | CHIKV | 33.8 | Pneumonia/Pneum onia |
| CHRF0110 | PCR (+) | m/0 | 10-Jul-17 | 60 (90)/150 | 16/Dis | CHIKV | 30.3 | Sepsis/Meningitis |

| Sample ID | DSH result | Age (m) | CSF collecti on date | WBC/ul (PMN%)/ Protein mg/dl | Hospital duration/ outcome | Organism detected in DSH | qPCR Ct | Provisional/ Final Diagnosis |
| --- | --- | --- | --- | --- | --- | --- | --- | --- |
| CHRF0111 | PCR (+) | m/8 | 18-Jul-17 | 42 (20)/35 | 7/Dis | CHIKV | 34.3 | AFC/Meningitis |
| CHRF0112 | PCR (+) | m/7 | 31-Aug-17 | 180 (10)/50 | 12/Dis | CHIKV | 29.3 | Meningitis/Meningitis |
| CHRF0113 | PCR (+) | m/1 | 18-Sep-17 | 90 (70)/150 | 17/Dis | CHIKV | 32.7 | Meningitis/Meningitis |
| CHRF0114 | Negative - Water | - | - | - | - | - | - | - |

ICSOL: Intracranial; UTI: Urinary tract infection; ARI: Acute respiratory tract infection; AGN: acute glomerulonephritis; AGE: acute gastroenteritis; AFC: Acute febrile convulsion; ASS: Acute stroke syndrome; NS: Nephrotic syndrome; LAMA: Left against medical advice; NA: the data are not available or applicable

**Table S4.** Case-based metagenomic data derived from all sequenced samples (n=115).

| Sample ID | mNGS output | Total reads | Nonhost reads | Non-host reads % | No of reads/million of the called pathogen | ERCC reads | RNA input |
| --- | --- | --- | --- | --- | --- | --- | --- |
| CHRF0052 | *E. coli* | 70250230 | 212176 | 0.3 | 465.9 | 41572416 | 17.2 |
| CHRF0064 | *S. pneumoniae* | 71227742 | 308236 | 0.4 | 3008.2 | 6063522 | 268.7 |
| CHRF0076 | *S. pneumoniae* | 63375670 | 98018 | 0.2 | 356.2 | 25658770 | 36.7 |
| CHRF0088 | *E. anophelis* | 85955422 | 162734 | 0.2 | 116.5 | 53048838 | 15.5 |
| CHRF0005 | *E. hormaechei* | 49785764 | 100862 | 0.2 | 465.9 | 23464986 | 28.0 |
| CHRF0017 | 0 | 44873536 | 71250 | 0.2 | - | 25545366 | 18.9 |
| CHRF0029 | *E. anophelis* | 54869610 | 373468 | 0.7 | 5758.7 | 46651102 | 4.4 |
| CHRF0041 | *K. pneumoniae* | 125053116 | 104932 | 0.1 | 21.0 | 101372810 | 5.8 |
| CHRF0001 | 0 | 102302082 | 216946 | 0.2 | - | 33395452 | 51.6 |
| CHRF0013 | 0 | 12772456 | 93332 | 0.7 | - | 11518604 | 2.7 |
| CHRF0025 | *S. pneumoniae* | 87017766 | 110194 | 0.1 | 36.0 | 15448776 | 115.8 |
| CHRF0037 | *S. pneumoniae* | 95449142 | 9212 | 0.0 | 7.5 | 516 | n/r |
| CHRF0049 | *S. pneumoniae* | 63460278 | 72664 | 0.1 | 18.8 | 37940022 | 16.8 |
| CHRF0061 | *S. pneumoniae* | 102545870 | 74388 | 0.1 | 74.2 | 25325942 | 76.2 |
| CHRF0073 | *S. pneumoniae* | 109138694 | 105100 | 0.1 | 554.8 | 6998288 | 364.9 |
| CHRF0085 | 0 | 80961046 | 158908 | 0.2 | - | 49273376 | 16.1 |
| CHRF0002 | *S. pneumoniae* | 141979356 | 2837444 | 2.0 | 19144.3 | 14875054 | 213.6 |
| CHRF0014 | *S. pneumoniae* | 5138438 | 1520 | 0.0 | 29.7 | 47076 | n/r |
| CHRF0026 | 0 | 150000000 | 216484 | 0.1 | 659.0 | 2209684 | 1670.2 |
| CHRF0038 | *S. pneumoniae* | 131599198 | 57492 | 0.0 | 15.4 | 3102744 | 1035.3 |
| CHRF0050 | 0 | 71994986 | 231934 | 0.3 | - | 55364152 | 7.5 |
| CHRF0062 | *S. pneumoniae* | 150000000 | 47112 | 0.0 | 40.9 | 4619612 | 786.7 |
| CHRF0074 | *S. pneumoniae* | 117315916 | 38716 | 0.0 | 5.1 | 1435378 | 2018.3 |
| CHRF0086 | *S. pneumoniae* | 66783636 | 69142 | 0.1 | 82.0 | 15896928 | 80.0 |
| CHRF0003 | *S. pneumoniae* | 56010100 | 88082 | 0.2 | 17.4 | 17328160 | 55.8 |
| CHRF0015 | 0 | 45328346 | 208836 | 0.5 | - | 42675752 | 1.6 |
| CHRF0027 | *H. influenzae* | 93564596 | 93914 | 0.1 | 56.3 | 43933980 | 28.2 |
| CHRF0039 | 0 | 88964960 | 47550 | 0.1 | - | 12434208 | 153.9 |
| CHRF0051 | *S. pneumoniae* | 150000000 | 42228 | 0.0 | 64.5 | 3295142 | 1113.2 |
| CHRF0063 | *N. meningitidis* | 83295482 | 338442 | 0.4 | 3.8, 3424.4 | 5436750 | 358.0 |
| CHRF0075 | *S. pneumoniae* | 111766028 | 12132 | 0.0 | 9.3 | 774910 | 3580.8 |
| CHRF0087 | *S. pneumoniae* | 69829522 | 30968 | 0.0 | 3.1 | 7217626 | 216.9 |
| CHRF0004 | 0 | 60813922 | 237838 | 0.4 | - | 45038786 | 8.8 |
| CHRF0016 | 0 | 56699154 | 127496 | 0.2 | - | 52922396 | 1.8 |
| CHRF0028 | 0 | 39764866 | 166378 | 0.4 | - | 37251668 | 1.7 |
| CHRF0040 | 0 | 85263302 | 282996 | 0.3 | - | 74509638 | 3.6 |
| CHRF0053 | 0 | 46124002 | 83482 | 0.2 | - | 44273832 | 1.0 |
| CHRF0065 | 0 | 44707958 | 182912 | 0.4 | - | 41350484 | 2.0 |
| CHRF0077 | 0 | 75937524 | 102300 | 0.1 | - | 66630914 | 3.5 |
| CHRF0089 | 0 | 6302354 | 184446 | 2.9 | - | 56770 | n/r |
| CHRF0006 | 0 | 80267448 | 145860 | 0.2 | - | 77153088 | 1.0 |
| CHRF0018 | 0 | 47931332 | 84642 | 0.2 | - | 44741152 | 1.8 |
| CHRF0030 | 0 | 128845460 | 113374 | 0.1 | - | 120459392 | 1.7 |
| CHRF0042 | 0 | 56269302 | 88386 | 0.2 | - | 53806038 | 1.1 |
| CHRF0054 | 0 | 109272300 | 127002 | 0.1 | - | 103557262 | 1.4 |
| CHRF0066 | 0 | 57896582 | 95220 | 0.2 | - | 54678742 | 1.5 |
| CHRF0078 | 0 | 42345140 | 63466 | 0.1 | - | 39235528 | 2.0 |
| CHRF0090 | 0 | 67481750 | 106452 | 0.2 | - | 62447376 | 2.0 |
| CHRF0007 | 0 | 52481998 | 126918 | 0.2 | - | 49759666 | 1.4 |
| CHRF0019 | 0 | 9003804 | 46672 | 0.5 | - | 8476152 | 1.6 |
| CHRF0031 | 0 | 65032886 | 190336 | 0.3 | - | 62205370 | 1.1 |
| CHRF0043 | 0 | 58224438 | 311660 | 0.5 | - | 53771664 | 2.1 |
| CHRF0055 | 0 | 65419488 | 115050 | 0.2 | - | 61576086 | 1.6 |
| CHRF0067 | 0 | 66072270 | 158258 | 0.2 | - | 58759328 | 3.1 |
| CHRF0079 | 0 | 60221934 | 86664 | 0.1 | - | 52120452 | 3.9 |
| CHRF0091 | 0 | 68421536 | 123050 | 0.2 | - | 62355960 | 2.4 |
| CHRF0008 | 0 | 56664264 | 127858 | 0.2 | - | 47826772 | 4.6 |
| CHRF0020 | 0 | 75031264 | 545870 | 0.7 | - | 64841106 | 3.9 |
| CHRF0032 | 0 | 52942812 | 94000 | 0.2 | - | 50417636 | 1.3 |
| CHRF0044 | 0 | 74679170 | 126652 | 0.2 | - | 70394724 | 1.5 |
| CHRF0056 | 0 | 58958602 | 100272 | 0.2 | - | 52111098 | 3.3 |
| CHRF0068 | 0 | 58263836 | 81770 | 0.1 | - | 53534916 | 2.2 |
| CHRF0080 | 0 | 58480150 | 77640 | 0.1 | - | 48353780 | 5.2 |
| CHRF0092 | 0 | 51999596 | 83734 | 0.2 | - | 46960542 | 2.7 |
| CHRF0009 | 0 | 42981288 | 122776 | 0.3 | - | 39365830 | 2.3 |
| CHRF0021 | 0 | 38396862 | 145968 | 0.4 | - | 35594906 | 2.0 |
| CHRF0033 | 0 | 50925620 | 68302 | 0.1 | - | 45749064 | 2.8 |
| CHRF0045 | 0 | 50455298 | 80706 | 0.2 | - | 48657850 | 0.9 |
| CHRF0057 | 0 | 50024148 | 95218 | 0.2 | - | 46905102 | 1.7 |
| CHRF0069 | 0 | 45846480 | 59750 | 0.1 | - | 42479872 | 2.0 |
| CHRF0096 | 0 | 49342992 | 69600 | 0.1 | - | 46646414 | 1.4 |
| CHRF0000 | 0 | 135087088 | 156310 | 0.1 | - | 130150782 | 0.9 |
| CHRF0081 | 0 | 64453194 | 38766 | 0.1 | - | 31709728 | 25.8 |
| CHRF0093 | 0 | 49670048 | 99216 | 0.2 | - | 43264818 | 3.7 |
| CHRF0010 | *Enterovirus B* | 61327650 | 70042 | 0.1 | 9.3 | 22230004 | 44.0 |
| CHRF0022 | 0 | 118348946 | 181994 | 0.2 | - | 70802894 | 16.8 |
| CHRF0034 | 0 | 60169132 | 112136 | 0.2 | - | 13310622 | 88.0 |
| CHRF0046 | 0 | 42195836 | 158576 | 0.4 | - | 25483100 | 16.4 |
| CHRF0058 | *M. tuberculosis* | 65072802 | 34314 | 0.1 | 9.5 | 3104784 | 499.0 |
| CHRF0070 | *B. cereus* | 69060574 | 98978 | 0.1 | 247.4 | 5797930 | 272.8 |
| CHRF0082 | *S. enterica* | 60637346 | 98414 | 0.2 | 97.8 | 19935498 | 51.0 |
| CHRF0094 | CHIKV | 61336096 | 412890 | 0.7 | 5495.9 | 28094424 | 29.6 |
| CHRF0011 | Mumps | 60553510 | 25850 | 0.0 | 0.7 | 7677666 | 172.2 |
| CHRF0023 | 0 | 53928028 | 285306 | 0.5 | - | 38094584 | 10.4 |
| CHRF0035 | Human herpes virus 6 (manual) | 96343858 | 16208 | 0.0 | - | 1643200 | 1440.8 |
| CHRF0047 | 0 | 57593626 | 54800 | 0.1 | - | 6349378 | 201.8 |
| CHRF0059 | *S. maltophilia* | 85986872 | 1362544 | 1.6 | 3240.9 | 38318362 | 31.1 |
| CHRF0071 | CHIKV | 58234020 | 85112 | 0.1 | 242.8 | 23316766 | 37.4 |
| CHRF0083 | 0 | 139233556 | 48796 | 0.0 | - | 286176 | n/r |
| CHRF0095 | 0 | 61324940 | 84592 | 0.1 | - | 19572240 | 53.3 |
| CHRF0012 | CHIKV | 8760674 | 56784 | 0.6 | 1830.1 | 3462238 | 38.3 |
| CHRF0024 | 0 | 98585738 | 86354 | 0.1 |  | 20945142 | 92.7 |
| CHRF0036 | Mumps (manual) | 72872708 | 62864 | 0.1 | 0 rpm, 2 total reads | 8675260 | 185.0 |
| CHRF0048 | 0 | 77389688 | 62590 | 0.1 | - | 58257176 | 8.2 |
| CHRF0060 | 0 | 81192516 | 193074 | 0.2 | - | 70471964 | 3.8 |
| CHRF0072 | 0 | 150000000 | 89218 | 0.1 | - | 31927140 | 92.1 |
| CHRF0084 | 0 | 59849024 | 57728 | 0.1 | - | 49233396 | 5.4 |
| CHRF0097 | CHIKV | 150000000 | 120342 | 0.1 | 24.9 | 130431532 | 3.8 |
| CHRF0098 | CHIKV | 150000000 | 187842 | 0.1 | 20.9 | 141728870 | 1.5 |
| CHRF0099 | CHIKV | 150000000 | 414592 | 0.3 | 6423.7 | 136277954 | 2.5 |
| CHRF0100 | CHIKV | 150000000 | 311012 | 0.2 | 8.0 | 95075520 | 14.4 |
| CHRF0101 | CHIKV | 150000000 | 1632694 | 1.1 | 20200.8 | 96521140 | 13.9 |
| CHRF0102 | CHIKV | 150000000 | 436436 | 0.3 | 111.5 | 113727008 | 8.0 |
| CHRF0103 | CHIKV | 150000000 | 381190 | 0.3 | 3254.2 | 121280236 | 5.9 |
| CHRF0104 | CHIKV | 150000000 | 371278 | 0.2 | 106.0 | 116330080 | 7.2 |
| CHRF0105 | CHIKV | 150000000 | 228930 | 0.2 | 881.3 | 105689416 | 10.5 |
| CHRF0106 | CHIKV | 150000000 | 359446 | 0.2 | 4169.4 | 117445236 | 6.9 |
| CHRF0107 | CHIKV | 150000000 | 171530 | 0.1 | 966.1 | 138854562 | 2.0 |
| CHRF0108 | CHIKV | 150000000 | 241328 | 0.2 | 248.9 | 131571546 | 3.5 |
| CHRF0109 | CHIKV | 150000000 | 157336 | 0.1 | 565.6 | 138399578 | 2.09 |
| CHRF0110 | CHIKV | 150000000 | 6795359 | 4.5 | 37386.1 | 16650300 | 200.3 |
| CHRF0111 | CHIKV | 150000000 | 223136 | 0.1 | 1065.6 | 136500544 | 2.5 |
| CHRF0112 | CHIKV | 150000000 | 25772 | 0.0 | 3.0 | 319182 | n/r |
| CHRF0113 | CHIKV | 150000000 | 1357528 | 0.9 | 34.8 | 72993550 | 26.7 |
| CHRF0114 | 0 | 150000000 | 231476 | 0.2 | - | 139254112 | 1.9 |

n/r: the RNA input could not be back-calculated from ERCC counts.

**Table S5.** List of microbes included in the logistic regression model as potential pathogens.

| **Bacteria** | **Viruses** |
| --- | --- |
| *Escherichia coli* | Enterovirus |
| *Shigella spp* | Adenovirus |
| *Ureaplasma spp* | Influenza A |
| *Streptococcus pyogenes* | Influenza B |
| *Salmonella spp* | Parainfluenza-1 |
| *Pseudomonas aeruginosa* | Parainfluenza-2 |
| *Staphylococcus aureus* | Parainfluenza-3 |
| *Streptococcus pneumoniae* | Respiratory syncytial virus |
| *Streptococcus agalactiae* | Parechovirus |
| *Klebsiella spp* | Enterovirus |
| *Haemophilus influenzae* | Human metapneumovirus |
| *Neisseria meningitidis* | Rubulavirus |
| *Mycoplasma pneumoniae* | Rhinovirus |
| *Chlamydia trachomatis* | Cytomegalovirus |
| *Chlamydophila pneumoniae* | Mumps virus |
| *Bordetella pertussis* | Chikungunya virus |
| *Stenotrophomonas spp* | Enterobacter |
| *Acinetobacter spp* | Alphavirus |
| *Mycobacterium tuberculosis* | Human herpesvirus |
| *Bacillus cereus* |  |
| *Elizabethkingia spp* |  |

### Method S1. Bioinformatic analysis and pathogen identification

Microbial pathogens were identified from raw sequencing reads using the IDseq Portal (https://idseq.net), a cloud-based, open-source bioinformatics platform designed for detection of microbes from metagenomic data (eFigure 1). IDseq scripts and user instructions are available at https://github.com/chanzuckerberg/idseq-dag and the graphical user interface web application for sample upload is available at https://github.com/chanzuckerberg/idseq-web. IDseq is conceptually based on previously implemented platforms,^1–3^ but is optimized for scalable Amazon Web Services (AWS) cloud deployment. Bioinformatics data processing jobs are carried out on demand as Docker containers using AWS Batch. Alignments to the National Center for Biotechnology Information (NCBI) database are executed on dedicated auto scaling groups (ASG) of Amazon Elastic Compute Cloud (EC2) instances, with the number of server instances varied with job load. Fast downloads of the NCBI database from the Amazon Simple Storage Service to each new server instance are enabled by the open-source tool s3mi (https://github.com/chanzuckerberg/s3mi). Initial alignment and removal of reads derived from the human genome is performed using the Spliced Transcripts Alignment to a Reference (STAR) algorithm.^4^ Low-quality reads, duplicates, and low-complexity reads are then removed using the Paired-Read Iterative Contig Extension (PRICE) computational package,^5^ the CD-HIT-DUP tool^6^ and a filter based on the Lempel-Ziv-Welch (LZW) compression score, respectively. A second round of human read filtering is carried out using bowtie2^7^ Remaining reads are queried against the most recent version of the NCBI nucleotide (NT) and non-redundant (NR) protein databases (updated monthly) using GSNAPL and RAPSearch2 respectively.^8,9^ Reads matching GenBank records in the superphylum Deuterostomia are removed, given the high likelihood that such residual reads are of human origin. The relative abundance of microbial taxa is calculated based on reads per million (rpM) mapped at the genus level. An overview of this pipeline is represented in eFigure 1.

## Method S2. Phylogenetic analysis of Chikungunya virus strains responsible for the meningitis outbreak in Bangladesh, 2017

To study the phylogeny of the Chikungunya virus (CHIKV), BLASTn was used to extract all complete CHIKV genomes that had greater then 85% identity to the draft genomes assembled using SPAdes v3.11.1.^1^ These genomes were then aligned using the default settings in MUSCLEv3.8.1551.^2^ Annotation from one of the NCBI genomes (accession number: HM045823) was used to divide the genome into coding, and non-coding regions, and ModelTest-NGv0.1.5 was used to identify the best-fitting evolutionary models for each genomic region. Using the best-fitting models for evolution for each genomic region, we reconstructed a maximum-likelihood phylogeny using RAxML-ng v0.6.0 using default settings.^3^ The maximum likelihood phylogeny was used to select a lineage of most closely related NCBI sequences to the assembled chikungunya virus genomes. We then created a time-resolved phylogeny for this lineage in BEAST v1.10.1 using substitution models selected using ModelTest-NG a strict molecular clock for all genomic sites, and selected the Bayesian skyline model as the tree prior.^4,5^ Two separate BEAST runs of 300 million Markov chain Monte Carlo iterations sampling every 50,000 iterations were used to explore the posterior distribution of phylogenies and evolutionary parameters. Convergence was visually checked after removing the first 50% of samples, and samples from the two chains were merged. A maximum clade credibility phylogeny was created using median node heights. Using the above described method, another time-resolved phylogeny of the Bangladesh CHIKV was generated using the genomes assembled earlier and E1 viral structural glycoprotein sequence from 2016-2017 South Asian cluster [Pakistan 2016 CHIKV outbreak (accession number: MF774613— MF774619, MF74074—MF74081), Bangladesh 2017 outbreak (accession number: MG697262—MG697282), Australia (accession number: KY751908), Italy (accession number: MG049915) Hong-Kong (accession number: MF503628) and India (accession number: KY057363)].

Accession number of Enterovirus B, CHRF0010, is MK468615 of the CHIKV genomes are CHRF_0071: MK468608; CHRF_0103: MK468609; CHRF_0094: MK468610; CHRF_0099: MK468611; CHRF_0101: MK468612; CHRF_0106: MK468613; CHRF_0108: MK468614; CHRF_0110: MK468615; CHRF_0104: MK468616; CHRF_0105: MK468617; CHRF_0107: MK468618; CHRF_0109: MK468619; CHRF_0012: MK468620; CHRF_0111: MK468621; CHRF_0112: MK468622; CHRF_0098: MK468623; CHRF_0100: MK468624; CHRF_0097: MK468625; CHRF_0102: MK468626; CHRF_0113: MK468627; CHRF_0012: MK468628; CHRF_0012: MK468629

